# DRG afferents that mediate physiologic and pathologic mechanosensation from the distal colon

**DOI:** 10.1101/2022.11.27.518103

**Authors:** Rachel L. Wolfson, Amira Abdelaziz, Genelle Rankin, Sarah Kushner, Lijun Qi, Ofer Mazor, Seungwon Choi, Nikhil Sharma, David D. Ginty

## Abstract

The properties of dorsal root ganglia (DRG) neurons that innervate the distal colon are poorly defined, hindering our understanding of their roles in normal physiology and gastrointestinal disease. Here, we report genetically defined subsets of colon innervating DRG neurons with diverse morphologic and physiologic properties. Four colon innervating DRG neuron populations are mechanosensitive and exhibit distinct force thresholds to colon distension. The highest threshold population, selectively labeled using *Bmpr1b* genetic tools, is necessary and sufficient for behavioral responses to high colon distension, which is partly mediated by the mechanosensory ion channel Piezo2. This HTMR population mediates behavioral over-reactivity to colon distension caused by inflammation in a model of inflammatory bowel disease. Thus, like cutaneous mechanoreceptor populations, colon innervating DRG afferents exhibit distinct anatomical and physiological properties and tile force threshold space, and genetically defined colon innervating HTMRs mediate pathophysiological responses to colon distension revealing a target population for therapeutic intervention.

## Introduction

Colon mechanosensation is critical for water resorption, motility through the GI tract, defecation, and, under pathologic distension, pain perception^1–3^. Sensory neurons extrinsic to the gut that innervate the distal colon and mediate these responses are those with cell bodies within the dorsal root ganglia (DRG), the DRG afferents. In addition to innervating internal organs, DRG afferents provide sensory innervation to the skin, where they have been extensively studied^4^. Low threshold mechanoreceptors (LTMRs) respond to gentle, innocuous mechanical forces^5–10^, whereas high threshold mechanoreceptors (HTMRs) start to respond at higher mechanical forces, and sometimes, although not always, encode force into the noxious range^11,12^. Cutaneous DRG afferents can also subdivided by conduction velocity, rate of adaption to sustained skin indentation, morphology, and genetic diversity, which has been used to create mouse genetic tools to label and manipulate the subtypes^5–10,12–15^.

Colon innervating DRG afferents have been categorized based on *ex vivo* physiologic recordings, or anatomical or genetic analysis^16–29^. Specifically, colonic DRG afferents have been defined by their response to probing at low or high mechanical forces, mucosal stroking, and circumferential wall stretch^19,30^, and their conduction velocities are in the Aδ or C-fiber range^3,19,31–33^. Moreover, a range of morphologically distinct colon innervating DRG neuron types have been identified through anterograde tracing^25^. These include endings that wrap around myenteric cell bodies (intraganglionic varicose endings (IGVE)), endings that branch into the circular or longitudinal muscle layers (intramuscular arrays (IMA)), endings that terminate within the submucosa, extend through the mucosa, or associate with blood vessels^25^. A large portion of colon innervating DRG neurons express calcitonin gene related peptide alpha (CGRP), encoded by *Calca*^24,25^, and have thus been classified as peptidergic.

Peptidergic DRG afferents that innervate the skin have either Aδ- or C-fibers^34,35^ and can respond to capsaicin, mustard oil, pruritogens, and others chemicals^36^ or high force mechanical stimuli^11,37^. Responding to noxious stimuli, including supraphysiologic distension and inflammation, is an important and clinically relevant role of colonic DRG afferents. Intracolonic inflammation leads to both behavioral^38^ and physiologic hypersensitivity to colonic distension^38–40^, but the identity of DRG afferent subtypes that mediate this response is unknown.

Single cell sequencing has revealed a large transcriptional heterogeneity within the DRG, with at least fifteen transcriptionally distinct sensory neuron subtypes^13,27,28,41,42^, however the identity and functions of the DRG neuron subtypes that innervate the colon are unclear. Here, we used mouse molecular genetic approaches, morphological analyses, functional imaging, electrophysiology, and behavior to define the properties and functional diversity of colon innervating DRG afferents and their roles in physiologic and pathophysiologic mechanosensation.

## Results

### Colon innervating DRG afferents have distinct morphologic and physiologic properties

Key to understanding sensation of the colon in both physiologic and pathologic states is appreciating the morphological and physiological properties of colon-innervating sensory neuron subtypes and the functions they subserve. In addition to DRG afferents, which provide extrinsic sensory innervation mainly to the distal gut, the GI tract has an extensive intrinsic system, the enteric nervous system (ENS), as well as additional extrinsic sensory and pre-ganglionic parasympathetic innervation provided by the vagus nerve, mainly to the upper GI tract, pre-ganglionic parasympathetic (splanchnic) innervation to the lower GI tract, and sympathetic nervous system (SNS) effector neurons terminating throughout the GI tract^43^. To establish mouse genetic tools for interrogating extrinsic DRG sensory neuron innervation of the distal GI tract, we first characterized labeling of peripheral nervous system subdivisions in mouse lines commonly used to manipulate DRG afferents, such as those in which a recombinase is expressed under control of the *Advillin* gene^44,45^.

4-hydroxytamoxifen (4-HT) treatment of *Advillin-CreER*; *R26^LSL-tdTomato^* mice promoted Cre-mediated tdTomato reporter expression in DRG, ENS, and SNS neuronal populations regardless of the developmental time 4-HT was given (Figure S1A-H). ENS labeling was defined by tdTomato^+^ myenteric cell bodies, while SNS labeling was identified by tdTomato^+^ cell bodies in sympathetic ganglia. In the skin and colon, control staining with anti-beta tubulin III (TUJ1) labeled all neurons (extrinsic and intrinsic), TH staining labeled all sympathetic endings and TH^+^ DRG afferents, and CGRP staining labeled both CGRPα^+^ DRG afferents and CGRPβ*^+^* ENS neurons, as CGRP antibodies cannot distinguish between the two isoforms. Intersecting Flp recombinase expressed from the *Advillin* gene locus (*Advillin^Flp^*) with Cre recombinase expressed in cells caudal to cervical level 2 (*Cdx2-Cre*) and a dual-recombinase tdTomato line (*R26^FSF-LSL-tdTomato^*) labeled both the DRG and SNS and excluded the ENS^46^ (Figure S1I-J). *PIRT^Cre^*, another commonly used DRG line, also robustly labeled the ENS (Figure S1K). Therefore, to selectively label DRG afferents, while excluding ENS, SNS, and vagal neurons, we used the *Phox2B-Flp* mouse line^47^, in which Flp recombinase is expressed in the ENS, SNS, and nodose ganglia, but not in the DRG (Figure S1L-M). This intersectional Cre-on, Flp-off genetic labeling approach (*Advillin-CreER*; *Phox2B-Flp*; *R26^FRT-LSL-ChR2-YFP-FRT^*) expressed yellow fluorescent protein (YFP) in cells in which Cre recombinase was expressed under control of the *Advillin* gene, but not in cells in which Flp recombinase was expressed under the control of the *Phox2B* gene. Indeed, in these mice YFP expression was observed in virtually all DRG neurons but not in the sympathetic ganglia, ENS cell bodies, or vagal afferents, as evidenced by lack of labeling in the nucleus tractus solitarius (NTS) (Figure 1A and S1N-O). This strategy defined genetic tools and intersectional genetic strategies for labeling colon innervating DRG neurons and confirmed that DRG afferents densely innervate the distal colon: most DRG neurons terminate within the myenteric plexus level, with fewer fibers found within the submucosal plexus level of the ENS (Figure 1A).

**Figure 1:**
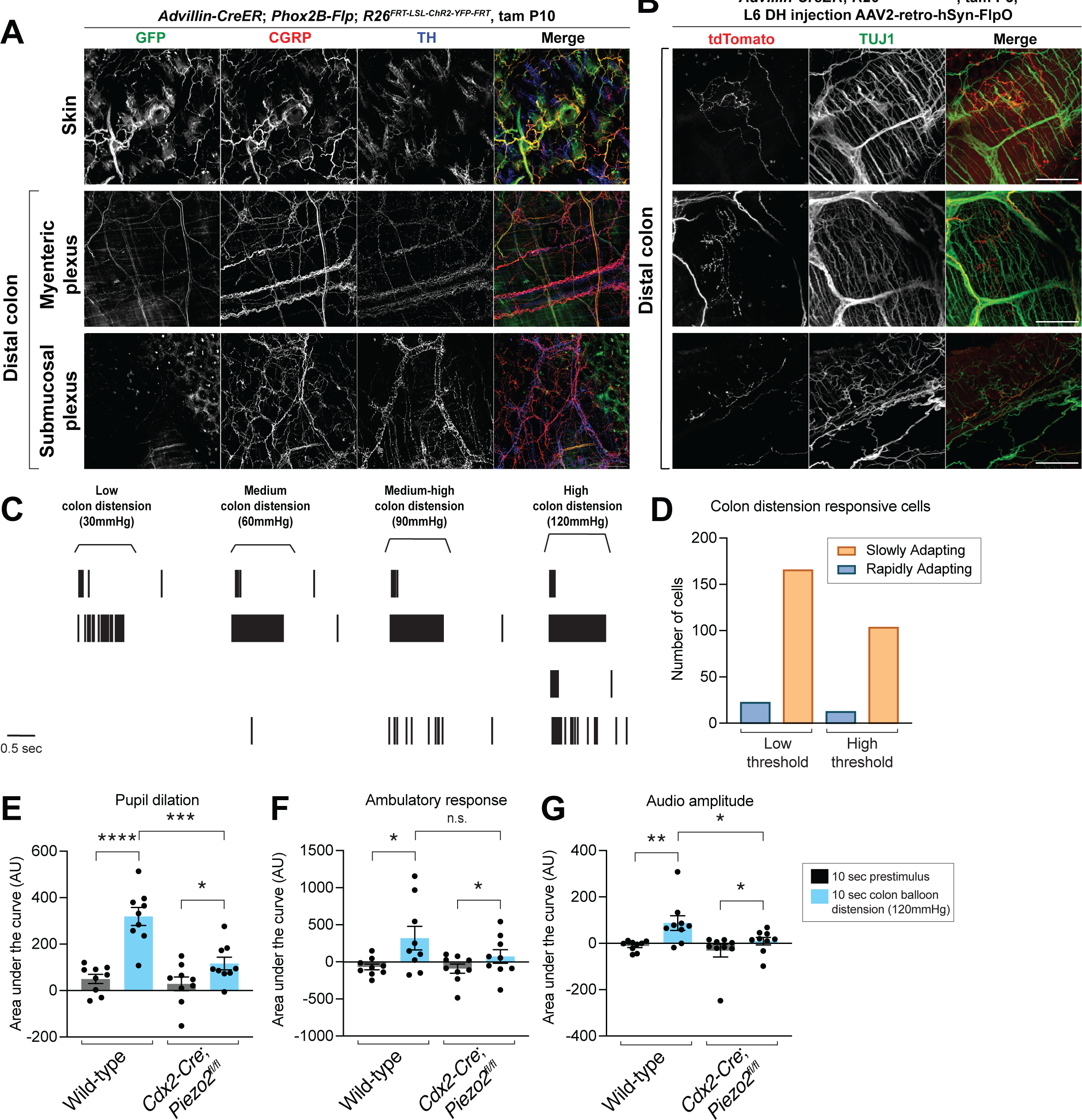
DRG afferents with distinct morphological and physiological properties innervate the distal colon and mediate Piezo2-dependent behavioral responses to colon distension. **A**: Whole mount immunostaining of skin and distal colon from *Advillin-CreER*; *Phox2B-Flp*; *R26^FRT-LSL-ChR2-YFP-FRT^* mice. Note that the GFP antibody labels YFP-expressing cells. **B**: AAV2-retro-hSyn-FlpO virus injection into the L6 dorsal horn (DH) of *Advillin-CreER*; *R26^FSF-LSL-tdTomato^* mice to sparsely label DRG afferent terminals in the distal colon. Tamoxifen time points are detailed in the methods unless otherwise specified in the figures. All scale bars are 100 μ unless otherwise specified. **C**: Example raster plots of single units from L6-S1 DRG multielectrode array (MEA) recordings. The top unit is classified as low-threshold rapidly adapting, followed by a low-threshold slowly adapting unit; the third unit is high-threshold rapidly adapting; the bottom unit is high-threshold slowly adapting. **D**: Quantification of the number of units in each category from L6-S1 DRG MEA recordings (data from N = 14 mice). **E-G**: The behavioral response to intracolonic high-threshold balloon distension in *Cdx2-Cre*; *Piezo2^fl/fl^* animals, measured by pupil dilation (**F**), movement (**G**), and vocalizations (**H**) (N = 9 mice per group; paired and unpaired t test for pupil dilation and accelerometer data; Mann-Whitney and Wilcoxon matched-pairs signed rank test for vocalization data). The following symbols are used in this and subsequent figures for P values: n.s., not significant; *, P < 0.05; **, P < 0.01; ***, P < 0.001; ****, P < 0.0001.

To assess the morphological diversity of DRG afferents that innervate the colon, we injected retro-AAV2-Flp virus into the L6-S1 dorsal horn of *Advillin-CreER*; *R26^FSF-LSL-tdTomato^* mice, such that tdTomato was expressed exclusively and sparsely in DRG afferents. The DRG afferent terminals in the colon exhibited multiple distinct morphologies, including IGVEs, IMAs, and endings that project through the mucosa towards the epithelium (Figure 1B). These ending morphologies reflect those that have been observed previously^25^.

We also assessed the diversity of physiological response properties of colon innervating DRG afferents. Colon innervating DRG afferents have distinct physiological responses in *ex vivo* preparations^20,24,48–50^, but the *in vivo* physiologic responses to colon distension have not been described. *In vivo* DRG multielectrode array (MEA) recordings revealed four categories of physiologic responses to colon distension: low-threshold rapidly adapting (RA); high-threshold RA; low-threshold slowly adapting (SA); and high-threshold SA (Figure 1C). The most common response types were SA, while the RA types were less common, although this may be due to a technical limitation of the assay, as RA neurons spike only a few times at the distension onset and/or offset and may therefore be more difficult to distinguish (Figure 1D).

We next assessed the role of Piezo2, the main mechanosensory ion channel in skin innervating DRG afferents^51^, in colon mechanosensation. To this end, we developed a behavioral assay to detect colon distension responses in awake animals. When distending an intra-colonic balloon to high force, mice exhibited a robust increase in pupil dilation as well as an increase in mobility and vocalization (Figure 1E-I). This may reflect a pain response, as a robust increase in pupil dilation, indicating a change in autonomic tone, movement, and vocalizations (including in the ultrasonic range) would be expected with a nociceptive stimulus. *Piezo2* ablation in cells caudal to cervical level 2 using *Cdx2-Cre*; *Piezo2^flox/flox^* mice, including DRG afferents but not enteric neurons (Figure S1I), decreased, but did not completely eliminate, the colon distension behavioral response. These findings indicate that Piezo2 is responsible for the majority of the behavioral response to high-threshold colon mechanosensation (Figure 1E-G).

### Five genetically-defined subtypes of DRG afferents innervate the distal colon

The finding that colon innervating DRG afferents exhibit a range of morphologies and physiologic colon distension responses implies an underlying afferent subtype heterogeneity. Therefore, to gain genetic access to colon innervating DRG neuron subtypes, we used tools that label a range of genetically defined DRG neurons, which were developed to explore the properties and functions of cutaneous DRG neuron populations. We focused on tools that label Aδ- and C-fiber DRG neuron subtypes because previous work has established that heavily δ myelinated neurons with Aβ conduction velocity do not innervate visceral organs^3,19,31–33^. We first used mouse intersectional genetic tools to selectively label MrgD^+^, TrkB^+^, and Tyrosine Hydroxylase (TH)^+^ DRG neurons^7,8,52,53^, which are well characterized to label cutaneous C-fiber HTMR, Aδ-LTMR, and C-LTMR DRG neuron populations, respectively. To selectively label MrgD^+^ DRG afferents while excluding the ENS, we employed the Cre-on, Flp-off strategy (*Mrgprd^CreER^*; *Phox2B-Flp*; *R26^FRT-LSL-tdTomato-FRT^*) and found that tdTomato-labeled MrgD^+^ neurons do not innervate the colon (Figure 2A, S2A). On the other hand, intersecting *TrkB^CreER^* and *Advillin^Flp^* mice was sufficient to label DRG neurons while excluding the SNS and ENS, and this approach revealed that a TrkB^+^ DRG neuron population densely innervates the distal colon (Figure 2B, S2B). To label TH^+^ DRG neurons while excluding SNS labeling, we used the Cre-on, Flp-off strategy with *TH^CreER^* and *Phox2B-Flp* (*TH^CreER^*; *Phox2B-Flp*; *R26^FRT-LSL-tdTomato-FRT^*) mice and found that tdTomato^+^ TH^+^ DRG afferents also densely innervate the distal colon (Figure 2C, S2C).

**Figure 2:**
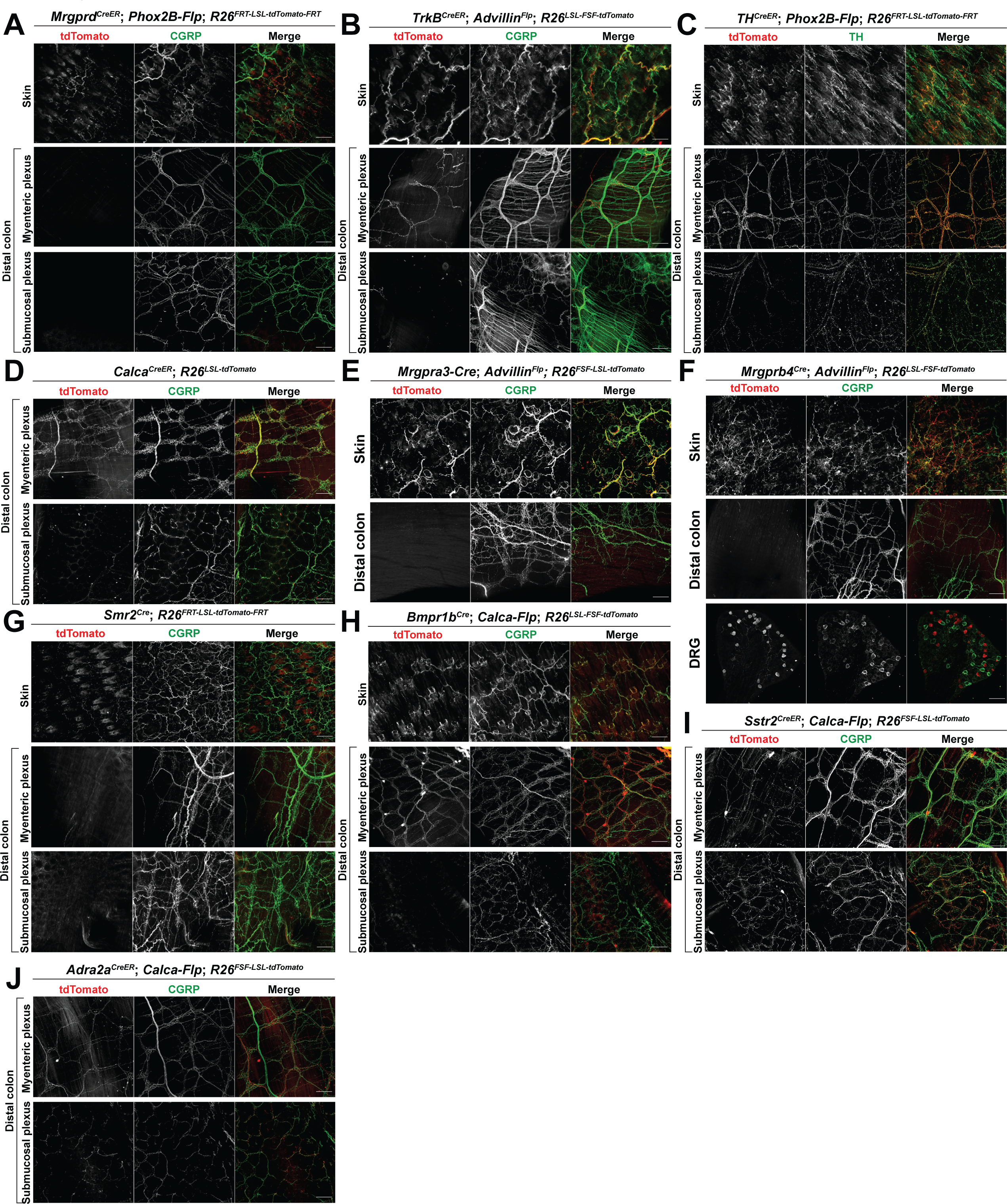
Five transcriptionally distinct DRG afferent subtypes that innervate the colon. Whole mount immunostaining in skin, distal colon, or DRGs from (**A**) Mrgprd^CreER^; Phox2B-_Flp; R26_FRT-LSL-tdTomato-FRT _mice; (**B**) TrkB_CreER_; Advillin_Flp_; R26_FSF-LSL-tdTomato _mice; (**C**) TH_CreER_;_ Phox2B-Flp; R26^FRT-LSL-tdTomato-FRT^ mice; (**D**) Calca^CreER^; R26^LSL-tdTomato^ mice; (**E**) Mrgpra3-Cre; Advillin^Flp^; R26^FSF-LSL-tdTomato^ mice; (**F**) Mrgprb4^Cre^; Advillin^Flp^; R26^FSF-LSL-tdTomato^ mice; (**G**) Smr2^Cre^; R26^FRT-LSL-tdTomato-FR^ mice ^T^; (**H**) Bmpr1b^Cre^; Calca-Flp; R26^FSF-LSL-tdTomato^ mice; (**I**) Sstr2^CreER^; Calca-Flp; R26^FSF-LSL-tdTomato^ mice; and (**J**) Adra2a^CreER^; Calca-Flp; R26^FSF-LSL-tdTomato^ mice. Tamoxifen time points are detailed in the methods unless otherwise specified in the figures. All scale bars are 100 μ unless otherwise specified.

We next focused on calcitonin gene-related peptide (CGRP) expressing DRG neuron subtypes, because CGRP^+^ DRG afferents are known to innervate the colon^25^. Multiple lines were used to characterize the CGRP population, two of which are BAC transgenics (*Calca-Flp* and *Calca-GFP*) and two others are *Calca* knock-ins (*Calca^CreER^*; *Calca^Lox-GFP-STOP-Lox-hDTR^*). These lines preferentially labeled CGRP^+^ DRG afferents with no SNS labeling and limited off-target ENS labeling (Figure 2D, S2D-F). The *Calca^CreER^*; *R26^LSL-tdTomato^* line labeled ∼2% of myenteric plexus cell bodies, indicating a small amount of ENS labeling (Figure S2G). Of the CGRP^+^ subsets of DRG neurons, as defined in Sharma et al.^13^, MrgA3^+^ neurons (and MrgB4^+^ neurons, which are a subset of the broader MrgA3^+^ population) and Smr2^+^ neurons (CGRP-ε population) do not innervate the GI tract (Figure 2E-G). Conversely, three subtypes of CGRP^+^ DRG afferents, defined by Bmpr1b (CGRP-η population), Sstr2 (CGRP-population), or Adra2a (CGRP-γ population) expression^13^, did innervate the colon (Figure 2H-J). For these three CGRP^+^ DRG neuron subsets, we used *Bmpr1b^Cre^* ^13^ and the newly developed *Sstr2^CreER^* and *Adra2a^CreER^* alleles and either intersection with *Calca-Flp* or the Cre-on, Flp-off strategy with *Phox2B-Flp* to confirm that the ENS and SNS were mostly excluded (Figure 2H-J, S2H-N). It is noteworthy that many genetically defined DRG neuron subtypes that innervate the colon had, as a population, the same general pattern observed when all DRG afferents were labeled, namely, dense innervation of the myenteric plexus but not the submucosal plexus, although the Sstr2^+^ and Adra2a^+^ CGRP subsets both had comparatively more innervation of the submucosal plexus. Thus, the TrkB^+^, TH^+^, Sstr2^+^ (CGRP-⍰), Adra2a^+^ (CGRP-γ), and Bmpr1b^+^ (CGRP-η) transcriptionally defined subtypes of DRG neurons have axons that innervate the distal colon, whereas MrgD^+^, MrgA3^+^, and Smr2^+^ (CGRP-ε) DRG neurons do not. Genetic access to five transcriptionally defined DRG subtypes enables physiological, morphological, and functional analysis of these populations.

### Genetic subtypes of DRG afferents have distinct morphologies in the distal colon

We and others have found that DRG afferents that innervate the distal colon exhibit a range of morphologies^25^. We hypothesized that each of the five transcriptionally defined DRG subtypes that innervate the colon exhibits a particular morphology and that collectively these subtypes account for much or all of the full range of DRG neuron morphologies. To test this, we performed sparse genetic labeling using mice harboring *TrkB^CreER^*, *TH^CreER^, Sstr2^CreER^*, *Adra2a^CreER^* alleles and a new *Bmpr1b^CreER^* allele (Figure S3A) and treated with a low dose of tamoxifen to label and visualize the endings of individual neurons for each of the five colon innervating DRG neuron subtypes. This analysis revealed that TrkB^+^ DRG afferents terminate exclusively in the myenteric plexus, with IGVE endings that wrap around ENS neuron cell bodies (Figure 3A, F-G). TH^+^ DRG neurons, on the other hand, either branched into the myenteric plexus with few myenteric wrappings or into the circular muscle layer, forming previously defined intramuscular array (IMA) endings (Figure 3B, F-G). Intersecting *Bmpr1b^CreER^* with *Calca-Flp* and a dual recombinase fluorescent reporter (*R26^FSF-LSL-tdTomato^*) allowed for sparse labeling of Bmpr1b^+^ DRG neurons (Figure S3B). As with TrkB^+^ DRG neurons, individual tdTomato labeled Bmpr1b^+^ neurons formed IGVEs, although with far fewer branches and wrappings around myenteric cell bodies; individual neurons typically wrapped around only 1-2 cell bodies (Figure 3C, F-G). tdTomato labeled Sstr2^+^ DRG neuron endings terminated in the myenteric plexus, submucosal plexus, or mucosa, in each case by branching but without any myenteric wrappings, consistent with previously described internodal strands (Figure 3D, F-G). tdTomato labeled Adra2a^+^ DRG neuron endings terminated either within the myenteric or submucosal plexuses, with few myenteric wrappings for those terminating in the myenteric plexus (Figure 3E-G). When comparing across the five different subtypes, TrkB^+^ endings were the most elaborate, with the longest length, most abundant branches, and most wrappings around myenteric cell bodies (Figure 3G-J). In summary, as observed for skin innervating DRG neurons subtypes, each of the five transcriptionally distinct, colon innervating DRG neuron subtypes exhibits a unique distal colon termination pattern and morphology, and collectively these colonic endings account for the range of morphologies observed in our random labeling experiments.

**Figure 3:**
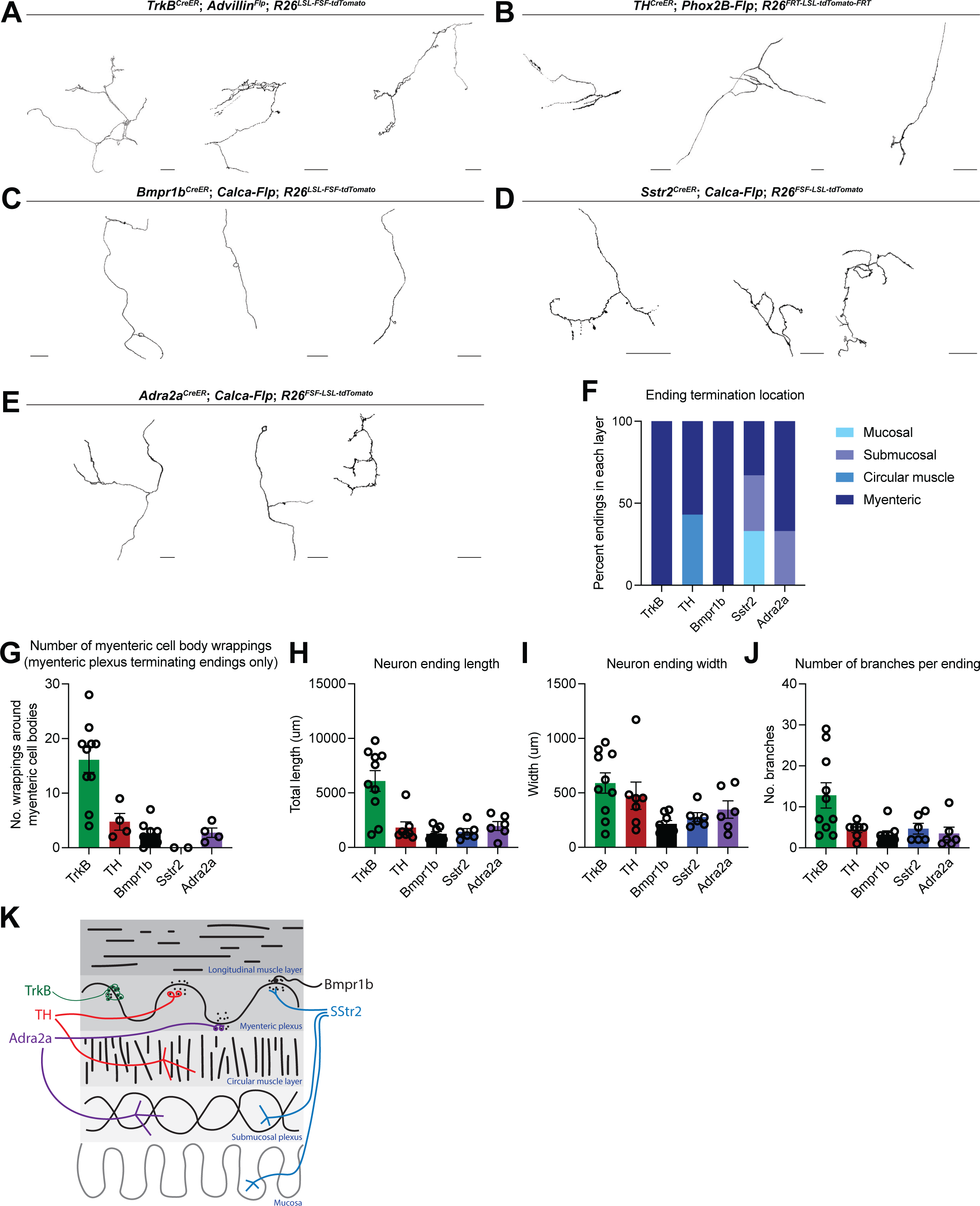
Five DRG neuron subtypes with distinct morphologies innervate the distal colon. **A-E**: Tracings of whole mount immunofluorescence from (**A**) TrkB^CreER^; Advillin^Flp^; R26^FSF-LSL-tdTomato^ mice; (**B**) TH^CreER^; Phox2B-Flp; R26^FRT-LSL-tdTomato-FRT^ mice; (**C**) Bmpr1b^CreER^; Calca-Flp; R26^FSF-LSL-tdTomato^ mice; (**D**) Sstr2^CreER^; Calca-Flp; R26^FSF-LSL-tdTomato^ mice; and (**E**) Adra2a^CreER^; Calca-Flp; R26^FSF-LSL-tdTomato^ mice, all treated with low amounts of tamoxifen to achieve sparse labeling. Tamoxifen time points are detailed in the methods unless otherwise specified in the figures. All scale bars are 100 μ unless otherwise specified. **F**: Graph summarizing ending termination location in the colon wall for the genetically labeled subtypes. **G**: Graph quantifying myenteric cell body wrappings observed with individual IGVEs by genetic subtype. Each point represents an individual ending. **H-J**: Graphs quantifying total ending length (**H**), width (**I**), or branches (**J**). Each point represents an individual ending. **K**: Diagram showing the colonic morphologies of distinct DRG afferent subtypes.

### At least four genetic subtypes of DRG afferents tile mechanical distension force space

Transcriptionally and morphologically distinct DRG neuron subtypes that innervate the skin have distinct physiological response properties. We therefore hypothesized that the transcriptionally and morphologically distinct colon innervating DRG neuron subtypes are also physiologically distinct. To determine which, if any, of the colon innervating DRG neuron subtypes respond to colon distension, we used the mouse genetic tools to express a GCaMP reporter selectively in each colon innervating subtype for *in vivo* L6-S1 DRG calcium imaging experiments (Figure 4A). Four of the five DRG neuron subtypes were assessed in this analysis; due to the sparse nature of labeling with the *Adra2a^CreER^* line, we were unable to perform functional experiments with this subtype. The TrkB^+^ DRG afferents responded to colon distension starting at low forces and in a RA manner (Figure 4B, S4A-B). Interestingly, the same colon responding TrkB^+^ neurons also often responded to back hairy skin pinch (Figure 4B). To ensure the colon responses were not due to slight movement of the skin during balloon distension of the colon, lidocaine was inserted into the colon. While colon distension responses were eliminated following lidocaine, skin pinch responses remained intact for the same neurons (Figure S4C-E). The TH^+^ DRG afferents responded to low-medium force colon distension in a SA manner (Figure 4C, S4F-G). It is noteworthy that colon innervating TH^+^ neurons were, on average, less sensitive than the TrkB^+^ neurons (Figure 4B, C, G).

**Figure 4:**
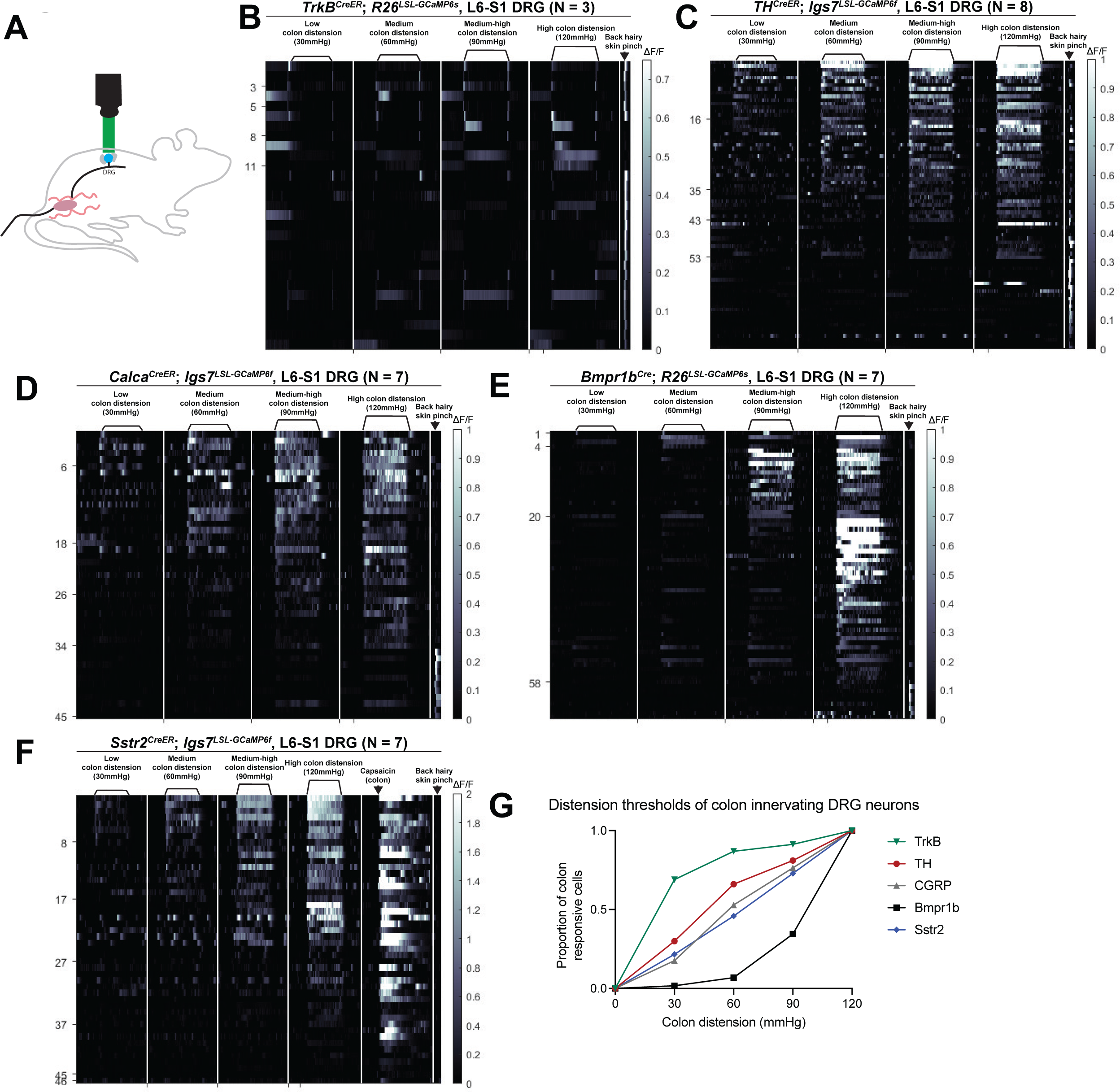
Four DRG neuron subtypes display distinct responses to colon distension and collectively tile force threshold space. **A**: A schematic for the DRG calcium imaging set up. **B-F**: *In vivo* DRG calcium imaging responses to colon distension in (**B**) *TrkB^CreER^*; *R26^LSL-GCaMP6s^* mice (N = 3); (**C**) *TH^CreER^*; *Igs7^LSL-GCaMP6f^* mice (N = 8); (**D**) *Calca^CreER^*; *Igs7^LSL-GCaMP6f^* mice (N = 7); (**E**) *Bmpr1b^Cre^*; *R26^LSL-GCaMP6s^* mice (N = 7); (**F**) and *Sstr2^CreER^*; *Igs7^LSL-GCaMP6f^* mice (N = 7). **G**: Graph depicting the distension thresholds of all colon innervating populations.

DRG calcium imaging from the entire CGRP^+^ population indicated that this transcriptionally and morphologically heterogenous population of DRG neurons respond mostly in a SA manner and to higher forces than either the TrkB^+^ or TH^+^ populations (Figure 4D, S4H-I). The Bmpr1b^+^ subtype of CGRP^+^ neurons also responded in a high-threshold SA manner (Figure 4E, S4J-K), although with a higher threshold than the CGRP population as a whole (Figure 4D, E, G). It is notable that the TrkB^+^, TH^+^, and Bmpr1b^+^ populations have force thresholds and physiologic response properties that are comparable to their genetic counterparts in the skin^7,8,11,13^. Finally, the Sstr2^+^ CGRP subtype, which expresses TrpV1, responded to medium-high force colon distension (Figure 4F, S4L-M). Consistent with their expression of TrpV1, intracolonic capsaicin activated the Sstr2^+^ afferents, but not Bmpr1b^+^ afferents, which do not express appreciable levels of TrpV1 (Figure 4F, S4N).

To complement the GCaMP imaging experiments, assess spike patterns, and estimate conduction velocities of the genetically labeled colon afferents, we next expressed an opsin in each distension-sensitive subtype and performed *in vivo* L6-S1 DRG electrophysiology using MEAs and optotagging with intracolonic light stimulation (Figure 5A-B). Consistent with the GCaMP imaging experiments, all opto-tagged TrkB^+^ units responded to the lowest colon distension forces and in a RA manner (Figure 5C-E, S5D). The optogenetic latency of this population was approximately 10ms and their waveform was narrow, consistent with A-fiber neurons (Figure 5D, F, S5A-C). Also consistent with the calcium imaging experiments, opto-tagged TH^+^ units responded to the lowest distension forces and in a SA manner (Figure 5C-E, S5D). In comparison to the TrkB^+^ neurons, TH^+^ neurons exhibited a longer optogenetic latency, which was in the C-fiber range, and a wider action potential waveform. Thus, like their cutaneous counterparts, colon innervating TrkB^+^ neurons are Aδ-LTMRs, whereas colon innervating TH^+^ neurons are C-LTMRs (Figure 5D, F, S5A-C).

**Figure 5:**
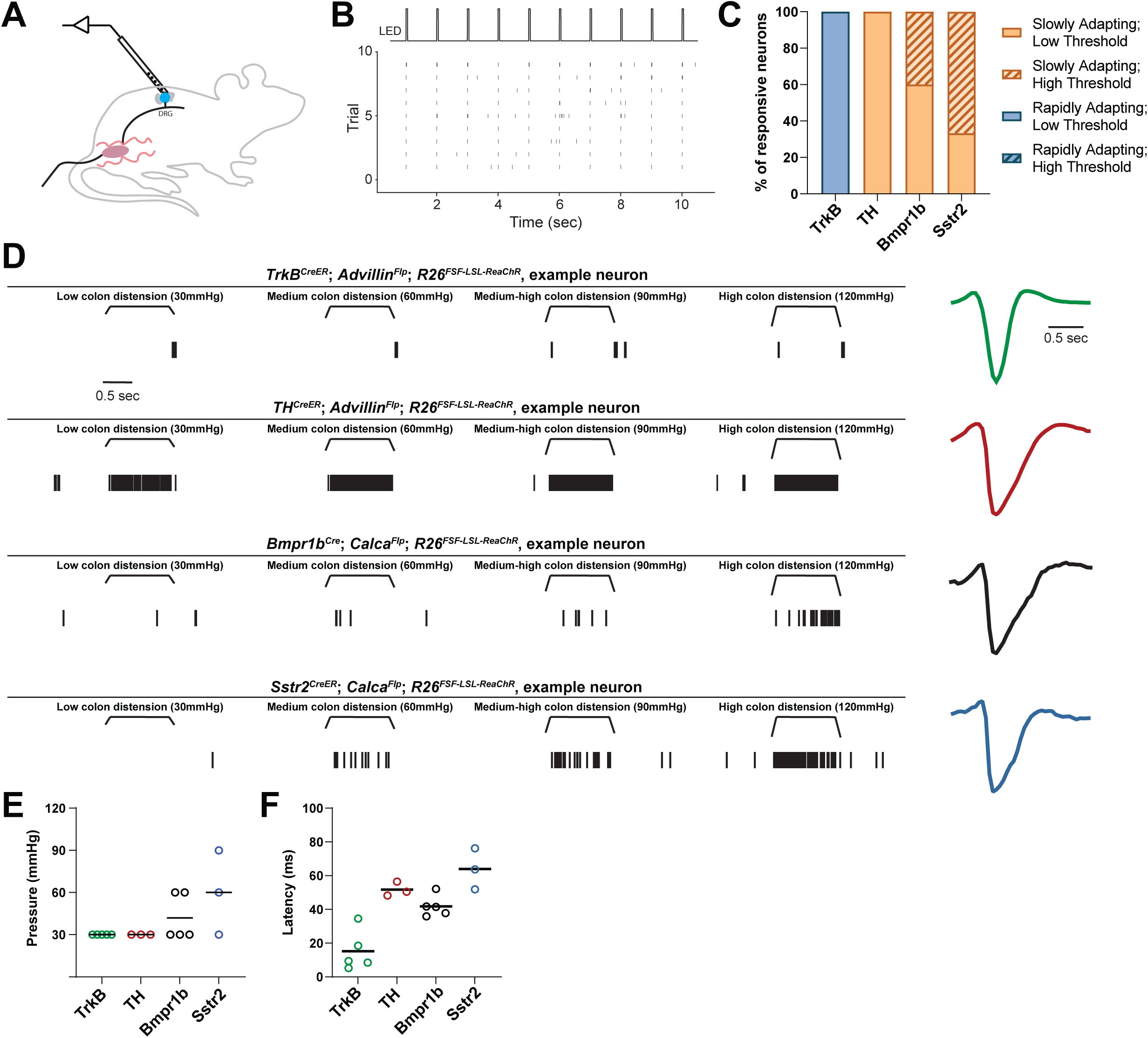
*In vivo* electrophysiological response properties of four colon innervating DRG neuron populations. **A**: A schematic of the DRG multielectrode array (MEA) set up. **B**: Example raster plot showing light-evoked responses of an opto-tagged neuron. **C**: Percent of neurons with different electrophysiological responses to colon distension in each genetic subtype. **D**: *In vivo* electrophysiological responses of one representative example neuron from each genetic class. Raster plot on the left; normalized extracellular action potential waveform on the right. **E**: Colonic distension pressure at which each neuron starts to respond in the MEA recording experiments by genetic subtype. Each point represents one neuron. **F**: Optogenetic latency by genetic subtype. Each point represents one neuron.

Analogous to the calcium imaging measurements, many opto-tagged Bmpr1b^+^ units started to respond to colon distension at higher forces and all responded in a SA manner (Figure 5C-E, S5D). This population had an optogenetic latency that was shorter than the TH^+^ C-LTMRs but longer than the TrkB^+^ Aδ-LTMRs, although with a wide waveform (Figure 5D, F, S5A-C). To further distinguish whether the Bmpr1b^+^ afferents were more likely A or C fibers, we performed a cell body size analysis and found that Bmpr1b^+^ cell bodies were considerably larger than Sstr2^+^ cell bodies, indicating that like their cutaneous counterparts, Bmpr1b^+^ afferents that innervate the colon are Aδ-HTMRs (Figure S5E). MEA recordings also confirmed that many Sstr2^+^ neurons begin to respond at higher thresholds, and all responded in a SA manner with long optogenetic latencies and wide action potential (AP) waveforms, which, together with the small cell body size, indicate that these are C-fiber neurons (C-HTMRs) (Figure 5C-F, S5A-E). Notably, both HTMR populations responded to lower distension forces in the MEA recordings compared with the calcium imaging experiments, which is likely due to the higher sensitivity of physiologic recordings. To determine the extent to which these four populations continue to encode distension forces into high force range, we performed a firing rate analysis of the MEA recording data. TrkB^+^ units do not appreciably increase their firing rate with increasing forces, consistent with their function as an LTMR (Figure S5F). Conversely, the Bmpr1b^+^ and Sstr2^+^ populations exhibited increased firing rates at higher forces, indicating that these populations continue to encode into the higher range, in line with their function as HTMRs, with the Bmpr1b^+^ displaying the highest total firing rates observed (Figure S5F). Interestingly, while robustly responding at the lowest threshold, the TH^+^ units increased their firing rate with increasing forces (Figure S5F). Taken together, the four genetic subpopulations of DRG afferents exhibit distinct physiological properties and responses to colon distension. While colon innervating TrkB^+^ neurons are Aδ-LTMRs and TH^+^ neurons are C-LTMRs, Bmpr1b^+^ and Sstr2^+^ neurons are Aδ-HTMR and C-HTMR populations, respectively. These four morphologically and physiologically distinct colon mechanoreceptor subtypes tile mechanosensory force threshold space, suggesting a population code to explain behavioral and perceptual responses to colon distension (Figure 4G).

### **Bmpr1b^+^ A**δ-HTMRs are necessary and sufficient for sensing high force colon distension

We next asked whether selective activation of any or all genetic subpopulations of DRG afferents is sufficient to mediate a behavioral response. To this end, we inserted an LED probe into the colon of mice expressing an opsin, either Channelrhodopsin-2 (ChR2) or red-shifted variant of Channelrhodopsin (ReaChR), in each of the four physiologically defined subtypes and recorded pupil dilation (indicative of a change in autonomic tone), ambulatory movement, and vocalizations in response to a light stimulus. In control experiments, light in the colon of wild-type animals did not lead to an appreciable response, while light on the skin of the back left hindpaw of wild-type mice led to a small amount of pupil constriction, presumably due to ambient light from the LED probe directly stimulating the retina, but no change in movement or vocalizations (Figure 6A-C and S6A-C).

**Figure 6:**
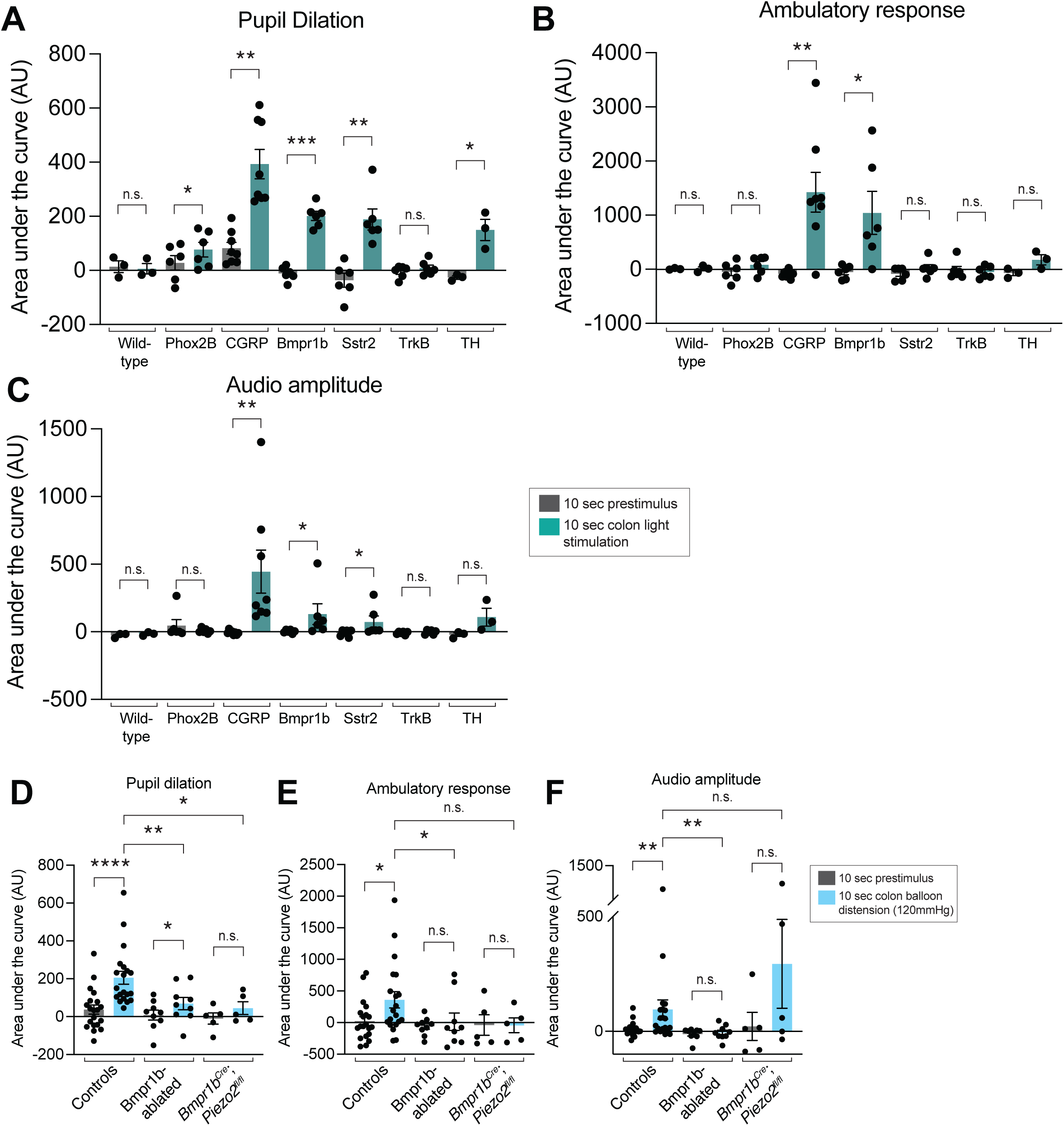
The Bmpr1b^+^ Aδ**-HTMR population is necessary and sufficient to drive a** behavioral response to colon stimulation. **A-C**: Behavioral responses (pupil dilation (**A**), movement (**B**), and vocalizations (**C**)) to colonic optogenetic stimulation of wild-type mice (N = 3), *Phox2B-Flp; R26^FSF-mCitrine-ReaChR^* mice (N = 6), *Calca-Flp*; *R26^FSF-ReaChR^* or *Calca^CreER^*; *R26^FRT-LSL-ChR2-YFP-FRT^* (N = 8), *Bmpr1b^Cre^*; *Calca-Flp*; *_R26_FSF-LSL-mCitrine-ReaChR* _mice (N = 6), *Sstr2*_*CreER*_; *Calca-Flp*; *R26*_*FSF-LSL-mCitrine-ReaChR* _or *Sstr2*_*CreER*_; *R26*_*LSL-mCitrine-ReaChR* _mice (N = 6), *TrkB*_*CreER*_; *Advillin*_*Flp*_; *R26*_*FSF-LSL-mCitrine-ReaChR* _mice (N = 6), and_ *TH^CreER^*; *Phox2B-Flp; R26^FRT-LSL-ChR2-YFP-FRT^* mice (N = 3). All comparisons are paired t tests except for the vocalization data for the *Phox2B-Flp; R26^FSF-mCitrine-ReaChR^* mice, *Calca-Flp*; *_R26_FSF-ReaChR* _or *Calca*_*CreER*_; *R26*_*FRT-LSL-ChR2-YFP-FRT* _mice, *Bmpr1b*_*Cre*_; *Calca-Flp*; *R26*_*FSF-LSL-mCitrine-ReaChR* _mice, and *Sstr2*_*CreER*_; *Calca-Flp*; *R26*_*FSF-LSL-mCitrine-ReaChR* _or *Sstr2*_*CreER*_; *R26*_*LSL-mCitrine-ReaChR* mice, which were analyzed using the Wilcoxon matched-pairs signed rank test. **D-F**: Behavioral responses (pupil dilation (**D**), movement (**E**), and vocalizations (**F**)) to high force colon balloon distension in Bmpr1b-ablated or *Bmpr1b^Cre^*; *Piezo2^fl/fl^* animals, compared to controls. Each point represents the average of data from an individual animal. (N = 20 for controls; N = 9 for Bmpr1b-ablated; N =5 for *Bmpr1b^Cre^*; *Piezo2^fl/fl^*. Comparisons for pupil dilation data are Wilcoxon matched-pairs signed rank test for the control paired groups, t tests for the Bmpr1b-ablated and *Bmpr1b^Cre^*; *Piezo2^fl/fl^* paired groups, and Mann Whitney tests for unpaired groups. Comparisons for accelerometer and vocalization data are Wilcoxon matched-pairs signed rank test for paired groups and Mann Whitney tests for unpaired groups).

We found that colonic or skin light stimulation of animals expressing ReaChR in TrkB^+^ Aδ-LTMRs, using *TrkB^CreER^*; *Advillin^Flp^*; *R26^FSF-LSL-ReaChR-mCitrine^* mice, did not elicit any obvious behavioral response (Figure 6A-C and S6A-C). Colonic light stimulation in animals expressing ChR2 in the TH^+^ C-LTMRs, using *TH^CreER^*; *Phox2B-Flp*; *R26^FRT-LSL-ChR2-YFP-FRT^* mice, increased pupil dilation without affecting vocalizations or movement (Figure 6A-C). This was distinct from the skin light stimulation behavioral response in these animals, in which there was no increase in pupil dilation, movement, or vocalizations (Figure S6A-C). On the other hand, optogenetic-activation of CGRP^+^ DRG afferents, using *Calca^CreER^*; *R26^LSL-ChR2-YFP^* or *Calca-Flp*; *R26^FSF-ReaChR^* mice, led to a strong behavioral response, with significant increases in pupil dilation, movement, and vocalizations during light stimulation of both the colon (Figure 6A-C) and skin (Figure S6A-C). Increases in pupil dilation, movement, and vocalizations (often in the ultrasonic range) may together indicate a nocifensive response, consistent with the known function of CGRP^+^ DRG afferents in the skin.

We next tested the individual CGRP subsets. As with the entire CGRP^+^ afferent population, optical activation of the Bmpr1b^+^ Aδ-HTMR subset of CGRP neurons, from both the colon and skin of *Bmpr1b^Cre^*; *Calca-Flp*; *R26^FSF-LSL-mCitrine-ReaChR^* mice, was sufficient to drive increases in pupil dilation, movement, and vocalizations. This response was less pronounced than colon light stimulation of the entire CGRP^+^ population (Figure 6A-C and S6A-C), which includes both the Bmpr1b^+^ Aδ-HTMRs and Sstr2^+^ C-HTMRs. Conversely, selective activation of the Sstr2^+^ C-HTMR colon-innervating subtype, using *Sstr2^CreER^*; *Calca-Flp*; *R26^FSF-LSL-mCitrine-ReaChR^* or *Sstr2^CreER^*; *R26^LSL-mCitrine-ReaChR^* mice, led to significant increases in pupil dilation and vocalizations, but not movement, whereas skin activation led to increases in pupil dilation, vocalizations, and movement (Figure 6A-C and S6A-C). Notably, these responses were not due to activation of the SNS, ENS, or vagal afferents, as colon light activation in animals expressing ReaChR in these populations, but not in the DRG (using *Phox2B-Flp*; *R26^FSF-ReaChR^* mice), led to a slight increase in pupil dilation without a change in vocalizations or movement (Figure 6A-C). Thus, optogenetic activation of Bmpr1b^+^ Aδ-HTMRs, and to a lesser extent Sstr2^+^ C-HTMRs, evoked behavioral responses analogous to the high colon distension response (Figure 1E-G), whereas TrkB^+^ Aδ-LTMRs and TH^+^ C-LTMRs did not.

Since Bmpr1b^+^ Aδ-HTMRs respond to high force mechanical distension of the colon and they evoke robust behavioral responses when optogenetically activated, we next asked whether this population is necessary for responses to painful colonic distension. Bmpr1b^+^ Aδ-HTMRs were ablated by expressing human diphtheria toxin receptor (hDTR) in these neurons using *Bmpr1b^Cre^*; *Calca^Lox-GFP-STOP-Lox-hDTR^* mice and treating them with diphtheria toxin (DTX) (Figure S6D). Indeed, ablation of Bmpr1b^+^ Aδ-HTMRs significantly decreased the high colon distension evoked behavioral response, indicating that this population is necessary for sensing high colon distension (Figure 6D-F) and is therefore a crucial colon innervating HTMR population. We also asked whether Piezo2 within Bmpr1b^+^ Aδ-HTMRs mediates behavioral responses to high force distension using *Bmpr1b^Cre^*; *Piezo2^flox/flox^* mice in which *Piezo2* was deleted in all Bmpr1b^+^ cells (Figure S6E). Piezo2 loss significantly decreased the high colon distension behavioral response (Figure 6D-F). *Piezo2* deletion in the Bmpr1b^+^ population, however, did not eliminate the physiologic response to high threshold colon distension in DRG calcium imaging (Figure S6F-H). These findings imply that Piezo2 expressed in Bmpr1b^+^ Aδ-HTMRs contributes to but is not fully responsible for the high colon distension response, analogous to noxious, high-threshold mechanosensation in the skin^54^.

### Bmpr1b^+^ HTMRs mediate inflammation-induced hypersensitivity in the colon

We next explored the role of Bmpr1b^+^ Aδ-HTMRs in colon physiology and pathophysiology. Under physiological circumstances, it is thought that DRG afferents impact motility through the GI tract^55^. We measured distal colon motility and found that Bmpr1b^+^ Aδ-HTMR ablation significantly increased lower GI tract motility time, indicating these neurons contribute to motility under normal physiological conditions (Figure 7A). We also tested whether these Aδ-HTMRs mediate responses to other, non-mechanical stimuli. In an open field assay, ablation of the entire CGRP^+^ DRG neuron population did not significantly alter behavioral responses to colonic mustard oil, a TrpA1 agonist (Figure S7A-C). Thus, the Bmpr1b^+^ Aδ-HTMRs, which are a subset of the CGRP^+^ population and do not appreciably express TrpA1^13^, while important for distension responses and motility, do not mediate the colonic mustard oil response.

**Figure 7:**
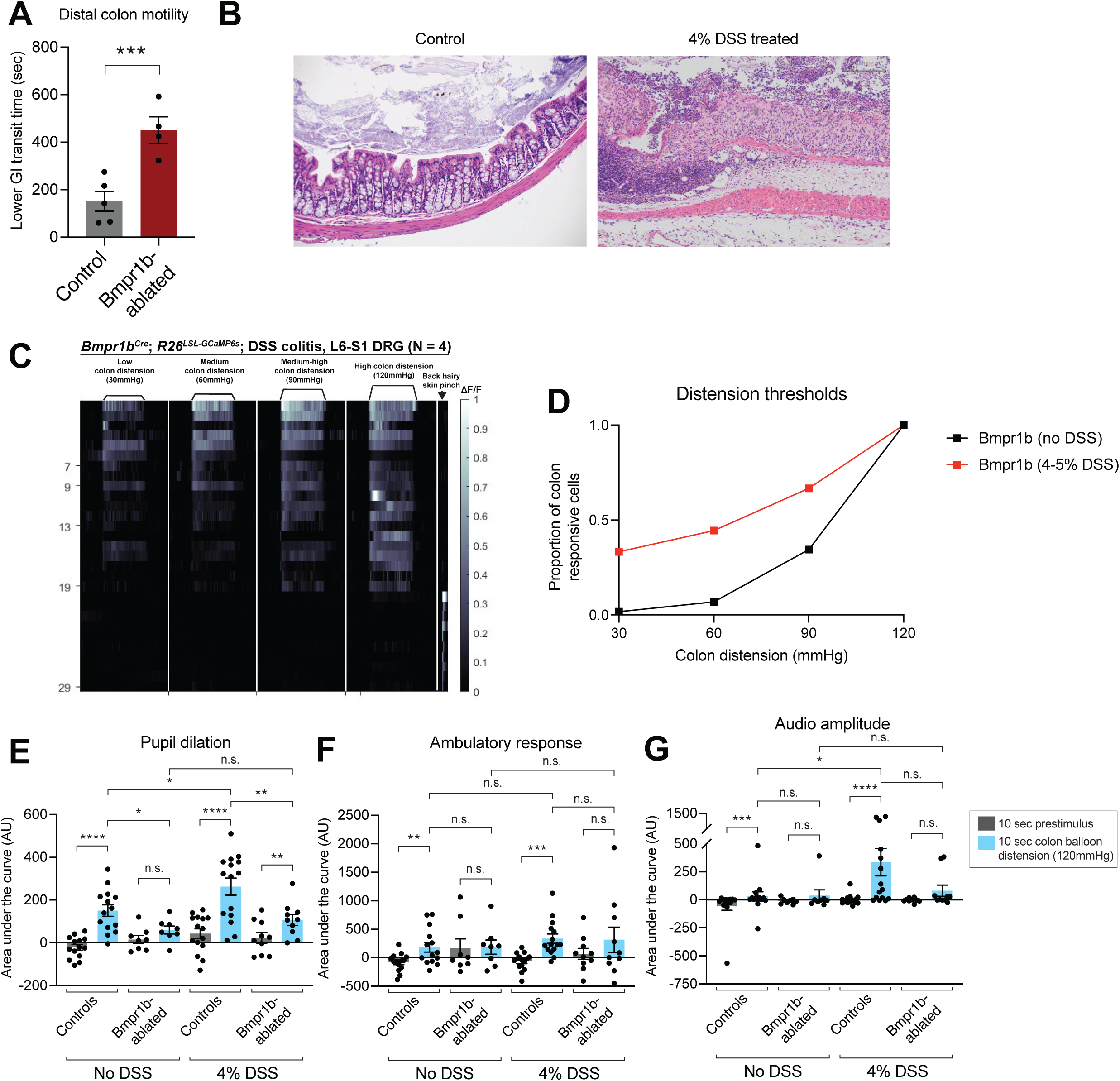
Bmpr1b^+^ Aδ**-HTMRs are necessary for colon physiology and inflammation-**induced colonic mechanical hypersensitivity. **A**: Distal colon motility in controls and Bmpr1b^+^ neuron-ablated animals. **B**: Colon hematoxylin and eosin (H&E) staining in an example control and DSS treated animal. Scale bar is 200 μm. **C**: *In vivo* DRG calcium imaging responses to colon distension in *Bmpr1b^Cre^*; *R26^LSL-GCaMP6s^* mice following DSS treatment (N = 4). **D**: Distension thresholds based on DRG calcium imaging of the Bmpr1b^+^ population in control or DSS treated animals. Data from *Bmpr1b^Cre^*; *R26^LSL-GCaMP6s^* (N = 7 for control; N = 4 for DSS treated). The control data are replotted from the data shown in Figure 4G. **E-G**: Behavioral responses (pupil dilation (**E**), movement (**F**), and vocalizations (**G**)) to high force colon balloon distension in Bmpr1b^+^ neuron-ablated mice or controls. Each point represents the average of data from an individual animal. (N = 14 control without DSS; N = 8 Bmpr1b-ablated without DSS; N = 15 control DSS-treated; N = 10 Bmpr1b-ablated DSS-treated. Comparisons for pupil dilation and accelerometer data are paired and unpaired t tests. Comparisons for vocalization data are Wilcoxon matched-pairs signed rank test for paired groups and Mann Whitney tests for unpaired groups.)

Previous reports indicate that colon inflammation causes behavioral over-reactivity and physiologic hypersensitivity to colon distension, though the identity of the DRG neuron population that mediates this response has remained elusive. Because Bmpr1b^+^ Aδ-HTMRs are both necessary and sufficient to mediate a nocifensive-like behavioral response, we tested their role in inflammation-induced hypersensitivity. Indeed, the Bmpr1b^+^ Aδ-HTMRs were more sensitive to distension in the setting of inflammation as compared with controls, based on calcium imaging experiments in *Bmpr1b^Cre^; R26^LSL-GCaMP6s^* DRGs after dextran sodium sulfate (DSS)-induced inflammation (Figure 7B-D). Furthermore, Bmpr1b^+^ Aδ-HTMR ablation (*Bmpr1b^Cre^*; *Calca^Lox-GFP-STOP-Lox-hDTR^* mice) reduced behavioral hypersensitivity to distension caused by DSS (Figure 7E-G, S7C). Control animals (untreated or DSS-treated) exhibited an increase in pupil dilation, movement, and vocalizations over their baseline in response to intracolonic balloon distension (Figure 7E-G). Ablated animals (untreated or DSS-treated) did not have an increased response over their baseline, apart from the DSS-treated Bmpr1b-ablated animals, which had an increase in pupil dilation only over their baseline, although this was significantly decreased in comparison to controls (Figure 7E-G). Additionally, while control animals displayed a significant increase in pupil dilation and vocalizations after DSS treatment, indicative of behavioral over-reactivity, Bmpr1b-ablated animals exhibited no change after DSS treatment (Figure 7E-G). Thus, Bmpr1b^+^ Aδ-HTMRs mediate pathophysiological responses to colon distension after chemical induced inflammation.

## Discussion

Colon mechanosensation is important in both GI physiology and pathophysiology. However, lack of genetic access to the colon innervating sensory neurons has hindered the understanding of the molecular and physiologic diversity of colon innervating neurons and their functions. Here, we labeled distinct transcriptionally defined subtypes of DRG afferents, while excluding the other sources of innervation to the GI tract, thus enabling selective genetic access to colon innervating sensory neuron subtypes. We found that the TrkB^+^ colon innervating DRGs form elaborate IGVEs, respond to low threshold colon distension in a RA manner with short optogenetic latencies, establishing their colonic function at Aδ-LTMRs. The TH^+^ population of colon innervating DRG neurons form either circular muscle IMAs or less elaborate IGVEs, respond to low force distension in a SA manner, and have longer optogenetic latencies, signifying their colonic function as C-LTMRs. We also established the properties of three CGRP^+^ subtypes in the colon. The Bmpr1b^+^ CGRP subtype, which form simple IGVEs with 1-2 myenteric wrappings, respond to high threshold colon distension in a SA manner, and have slightly faster optogenetic latencies than the C-LTMRs and large cell bodies, implying their function as Aδ-HTMRs. The Sstr2^+^ CGRP subtype form endings that branch into the myenteric plexus, submucosa, or mucosa, respond to high threshold colon distension and intracolonic capsaicin in a SA manner, and have optogenetic latencies that are slower than the C-LTMRs, establishing their function as C-HTMRs. Finally, Adra2a^+^ CGRP neurons form IGVEs with few myenteric wrappings or branch into the submucosal layer, however, due to the sparseness of labeling with the *Adra2a^CreER^* mouse line, their physiological properties remain elusive and will need to be determined in future studies.

As in the skin, the most widely studied peripheral sensory organ, both LTMRs and HTMRs innervate the colon. Moreover, in both the skin and colon, mechanosensory neurons exhibit a range of innervation patterns, morphologies, and force thresholds. Remarkably, there is a conservation of properties of three of the four subtypes that innervate the colon and skin. Thus, like cutaneous Aδ-LTMRs^7^, the TrkB^+^ neurons that innervate the colon are Aδ-LTMRs and respond in a RA-manner, whereas the TH^+^ afferents in the colon exhibit low force thresholds and slow conduction velocities in the C-fiber range, thus resembling C-LTMRs that innervate the skin^8^. Likewise, the Bmpr1b^+^ CGRP afferents are Aδ-HTMRs, similar to their cutaneous counterparts^11,13^. This conservation of response properties despite distinct end organ morphologies and environments in the skin and colon implies that the genetic composition of a mechanosensory neuron type dictates its mechanical force thresholds and adaptation to sustained stimuli.

Conversely, the TrpV1^+^ DRG afferents are thought to be mechanically insensitive but heat-responsive in the skin, as TrpV1 ablation leads to deficits in heat but not mechanical sensitivity^56^. Colon innervating, TrpV1^+^, Sstr2^+^ C-HTMRs, on the other hand, respond to both high-threshold colon distension as well as intracolonic capsaicin. Multiple possibilities may explain this dichotomy, including distinct microenvironments of the skin and colon, perhaps accounting for distinct responses to mechanical force, or genetic heterogeneity within the Sstr2-expressing population, which will be explored in future studies.

We further find that, like their cutaneous counterparts, most mechanosensitivity of colon innervating DRG afferents is mediated by Piezo2. However, some of the high-threshold force sensitivity of colon innervating DRG afferents is Piezo2-independent, implying that either a high-threshold mechanosensitive ion channel or tissue damage responses mediated by other signals, perhaps P2X receptors, account for at least some of the high force responses, also analogous to high threshold cutaneous responses to noxious touch.

Interestingly, Bmpr1b^+^ Aδ-HTMR ablation prolongs colon motility, implicating an important role of this HTMR population in physiologic function. It is notable that this DRG afferent’s role in the distal colon appears to be necessary to sense contents and increase expulsion, though future work will be needed to characterize its role in other parts of the GI tract.

Finally, while it is well-established that colon inflammation, such as in the setting of inflammatory bowel disease, leads to mechanical hypersensitivity, the genetic identity, morphologic and physiologic properties, and functions of intrinsic and extrinsic neurons responsible for this behavioral change have remained elusive. We found that Bmpr1b^+^ Aδ-HTMRs become more sensitive and mediate pathophysiological responses to colon distension after inflammation and may therefore represent a novel and clinically relevant therapeutic target for treating colon pain. Further work is needed to deeply probe the genetic expression profiles of Bmpr1b^+^ Aδ-HTMRs and other DRG afferent populations^13^ to uncover potential molecular targets to treat abdominal pain.

## Figure legends

## Supplementary Figures

**Supplementary Figure 1:**
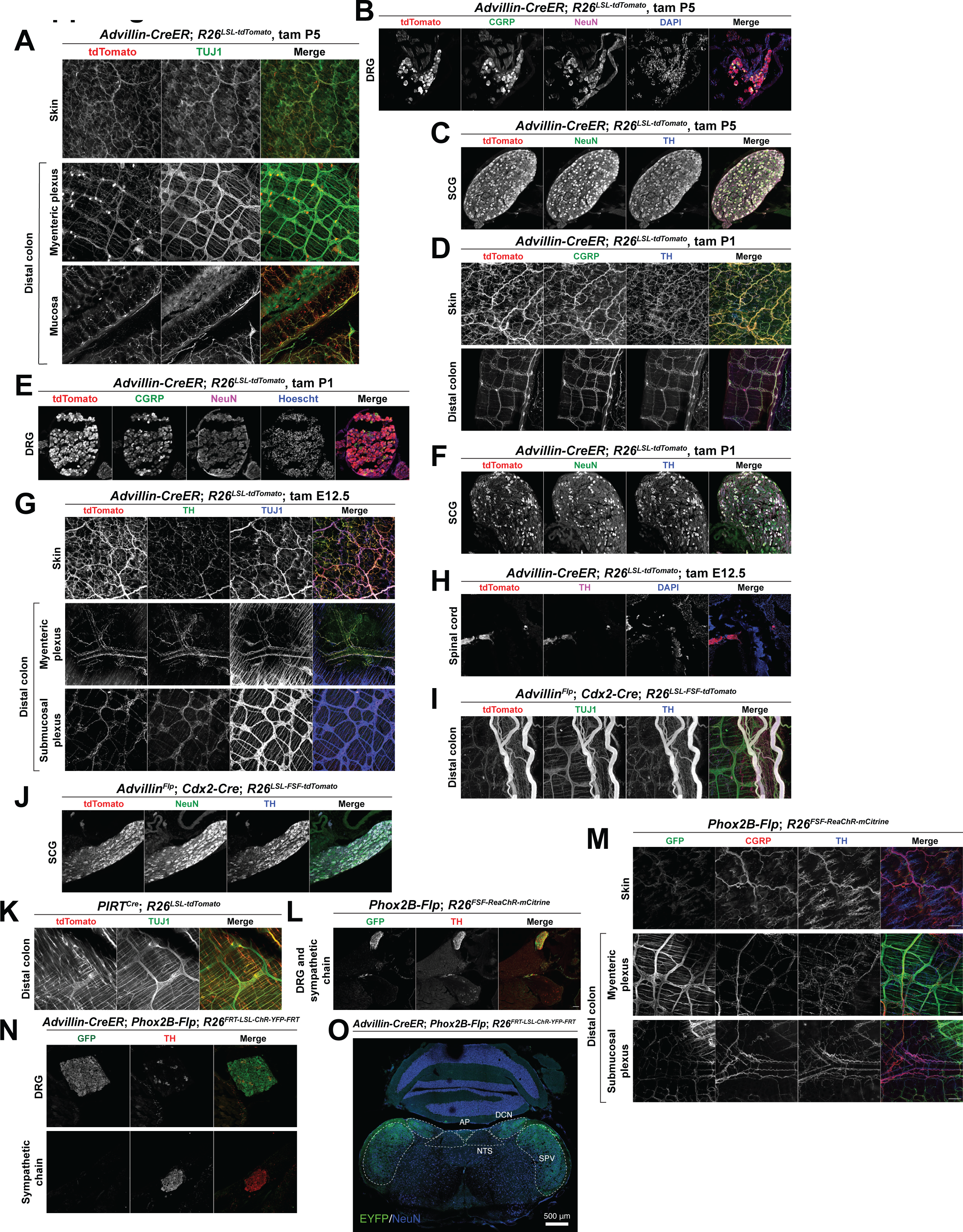
Peripheral nervous system labeling using mouse lines targeting the DRG. **A-M**: Skin and colon whole mount immunofluorescence or DRG, superior cervical ganglia (SCG), sympathetic chain, or spinal cord immunofluorescence in (**A-C**) *Advillin-CreER*; *R26^LSL-tdTomato^* mice treated with tamoxifen at P5; (**D-F**) *Advillin-CreER*; *R26^LSL-tdTomato^* mice treated with tamoxifen at P1; (**G-H**) *Advillin-CreER*; *R26^LSL-tdTomato^* mice treated with tamoxifen at E12.5; (**I-J**) *Advillin^Flp^*; *Cdx2-Cre*; *R26^FSF-LSL-tdTomato^*; (**J**) *PIRT^Cre^*; *R26^LSL-tdTomato^*; (**L-M**) *Phox2B-Flp*; *R26^FSF-ReaChR-mCitrine^*; or (**N**) *Advillin-CreER*; *Phox2B-Flp*; *R26^FRT-LSL-ChR2-YFP-FRT^* animals. Tamoxifen time points are detailed in the methods unless otherwise specified in the figures. All scale bars are 100 μm unless otherwise specified. **O**: Brainstem immunofluorescence in *Advillin-CreER*; *Phox2B-Flp*; *R26^FRT-LSL-ChR2-YFP-FRT^* (area postrema (AP), nucleus tractus solitarius (NTS), dorsal column nucleus (DCN), spinal trigeminal nucleus (SPV)). Scale bar is 500 μm.

**Supplementary Figure 2:**
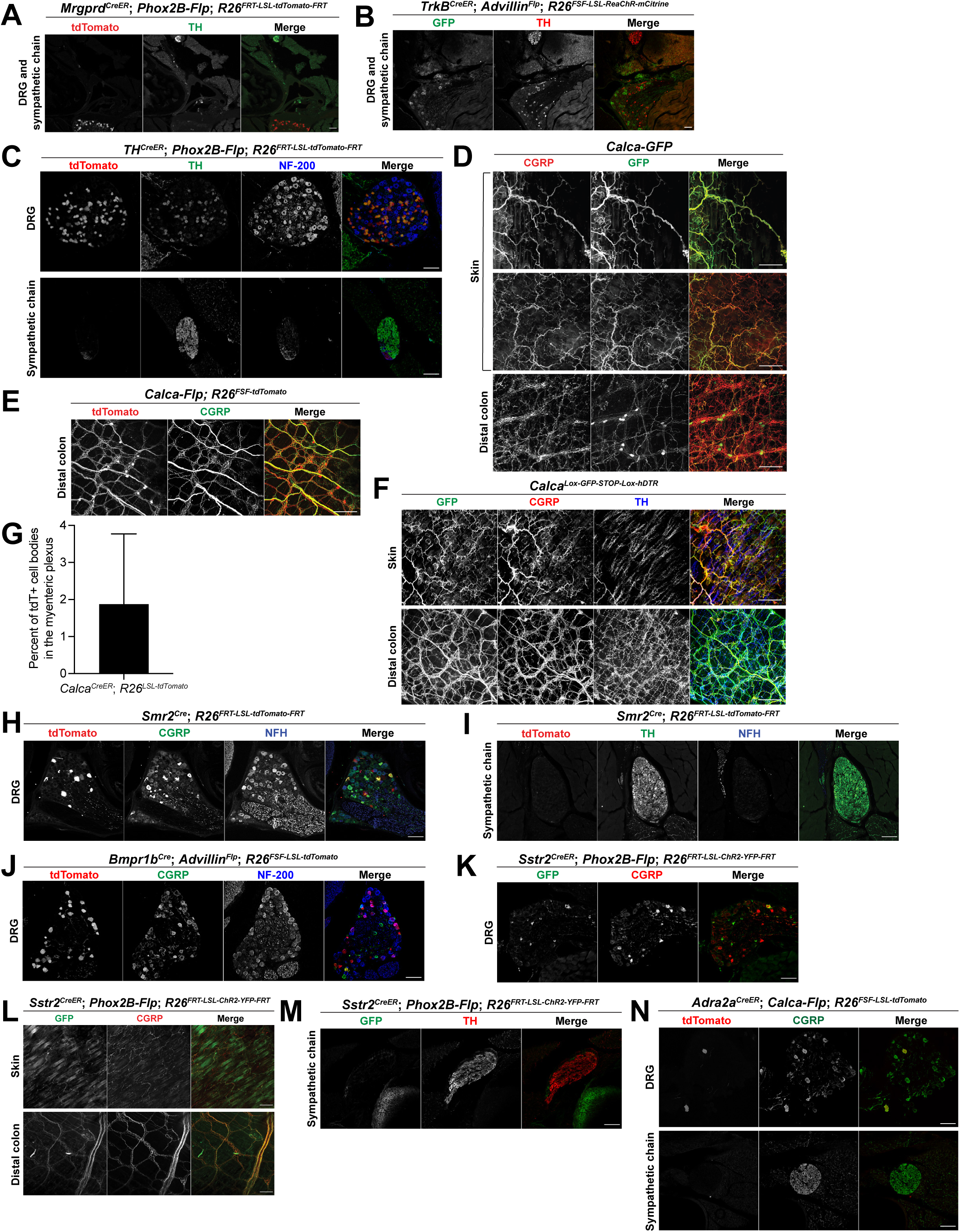
Peripheral nervous system labeling in mouse lines targeting distinct DRG neuron subtypes. **A-F**: Skin and colon whole mount immunofluorescence or DRG and sympathetic chain immunofluorescence in (**A**) *Mrgprd^CreER^*; *Phox2B-Flp*; *R26^FRT-LSL-tdTomato-FRT^* mice; (B) *TrkB^CreER^*; *_Advillin_Flp*_; *R26*_*FSF-LSL-ReaChR-mCitrine* _mice; (**C**) *TH*_*CreER*_; *Phox2B-Flp*; *R26*_*FRT-LSL-tdTomato-FRT* _mice;_ (**D**) *Calca-GFP*; (**E**) *Calca^Flp^*; *R26^FSF-tdTomato^*; (**F**) *Calca^Lox-GFP-STOP-Lox-hDTR^* mice. **G**: Percent of tdTomato labeled myenteric plexus cell bodies in colon whole mount immunofluorescence from *Calca^CreER^*; *R26^LSL-tdTomato^* mice. **H-N**: DRG and sympathetic chain immunofluorescence in (**H-I**) *Smr2^Cre^*; *R26^FRT-LSL-tdTomato-FRT^* mice; (**J**) *Bmpr1b^Cre^*; *Advillin^Flp^*; *R26^FSF-LSL-tdTomato^* mice; (**K-M**) *Sstr2^CreER^*; *Phox2B-Flp*; *R26^FRT-LSL-ChR2-YFP-FRT^* mice; and (**N**) *Adra2^CreER^*; *Calca-Flp*; *R26^FSF-LSL-tdTomato^* mice. Tamoxifen time points are detailed in the methods unless otherwise specified in the figures. All scale bars are 100μ m unless otherwise specified.

**Supplementary Figure 3:**
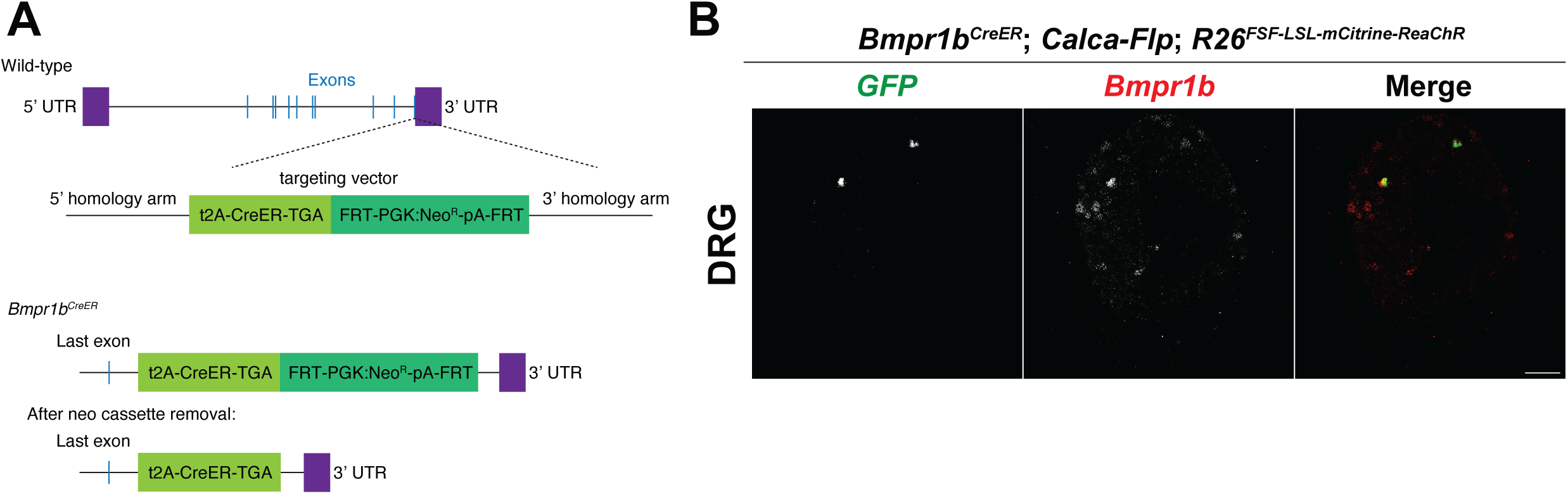
Generation of a mouse line to sparsely label Bmpr1b^+^ neurons. **A**: *Bmpr1b^CreER^* targeting vector schematic. **B**: DRG *in situ* hybridization in *Bmpr1b^CreER^*; *Calca-Flp*; *R26^FSF-LSL-mCitrine-ReaChR^* mice treated with 2 mg tamoxifen at P21 and sacrificed at least fourteen days later.

**Supplementary Figure 4:**
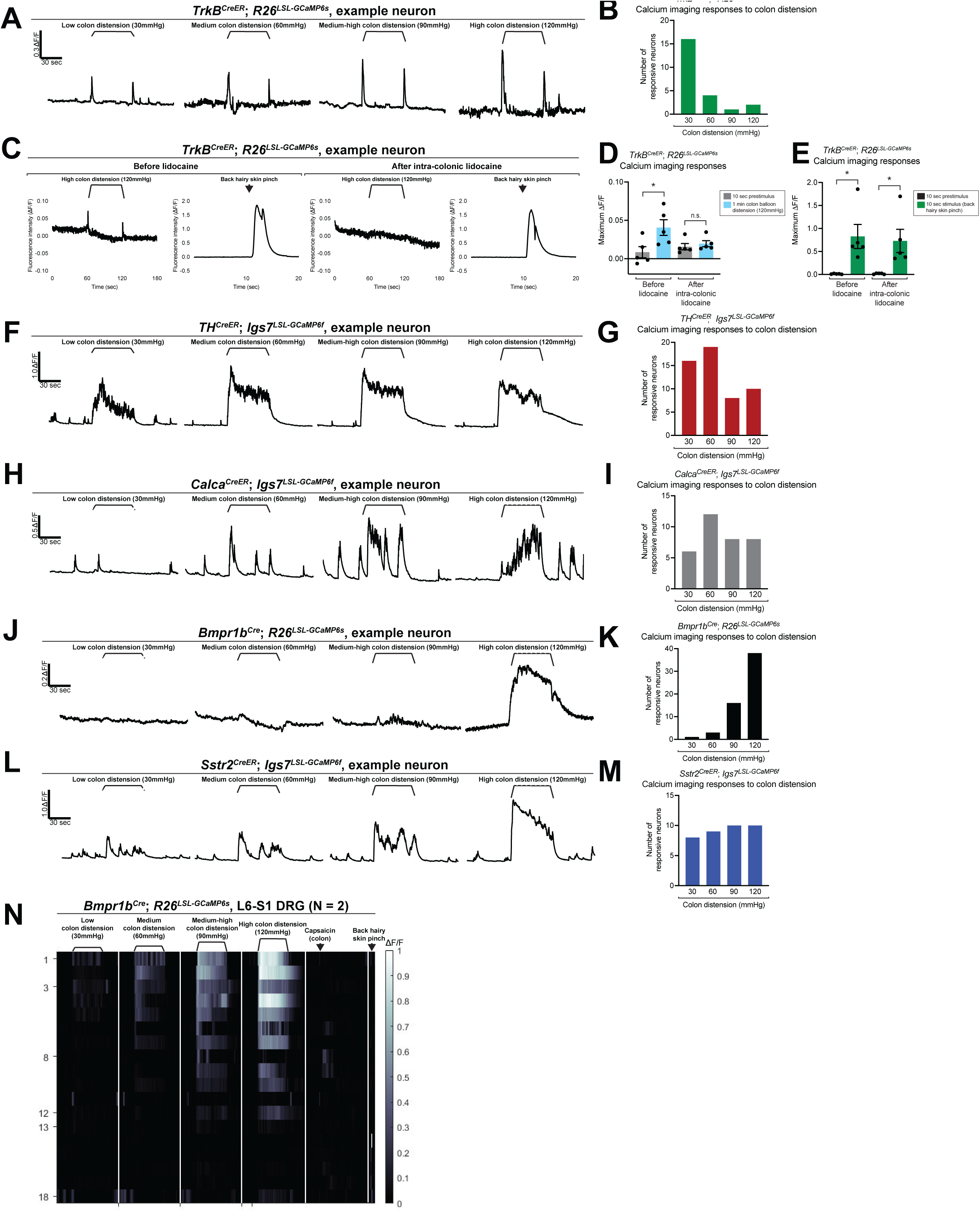
Four DRG afferent subtypes have distinct colonic distension responses based on *in vivo* calcium imaging. **A**: Calcium traces from an example TrkB^+^ neuron. **B**: Graph with distension thresholds of all TrkB^+^ responsive cells (data from N = 3 mice). **C**: Calcium traces showing response to colon distension or skin pinch in an example TrkB^+^ neuron before and after intra-colonic lidocaine. **D**: Maximum fluorescence intensity in the 10 seconds prior and 1 minute during intracolonic balloon distension (120mmHg) before or after intracolonic lidocaine in TrkB^+^ cells (each point respresents an individual neuron; N = 3 animals, 2 *TrkB^CreER^*; *R26^LSL-GCaMP6s^* mice and 1 *TrkB^CreER^*; *R26^LSL-GCaMP6s^*; *Piezo2^fl/+^* mouse). **E**: Maximum fluorescence intensity in the 10 seconds prior and during back hairy skin pinch before or after intracolonic lidocaine in TrkB^+^ neurons (each point represents an individual neuron; N = 3 animals, 2 *TrkB^CreER^*; *R26^LSL-GCaMP6s^* mice and 1 *TrkB^CreER^*; *R26^LSL-GCaMP6s^*; *Piezo2^fl/+^* mouse). **F**: Calcium traces depicting colon distension responses of an example TH^+^ neuron. **G**: Graph with distension thresholds of all TH^+^ responsive neurons recorded (data from N = 8 mice). **H**: Calcium traces from an example CGRP^+^ neuron. **I**: Graph with distension thresholds of all CGRP^+^ responsive neurons (data from N = 7 mice). **J**: Calcium traces from an example Bmpr1b^+^ neuron. **K**: Graph with distension thresholds of all Bmpr1b^+^ responsive neurons (data from N = 7 mice). **L**: Calcium traces from an example Sstr2^+^ neuron. **M**: Graph with distension thresholds of all Sstr2^+^ responsive neurons (data from N = 7 mice). **N**: *In vivo* DRG calcium imaging responses to colon distension and intracolonic capsaicin in Bmpr1b^Cre^; R26^LSL-GCaMP6s^ (N = 2) animals.

**Supplementary Figure 5:**
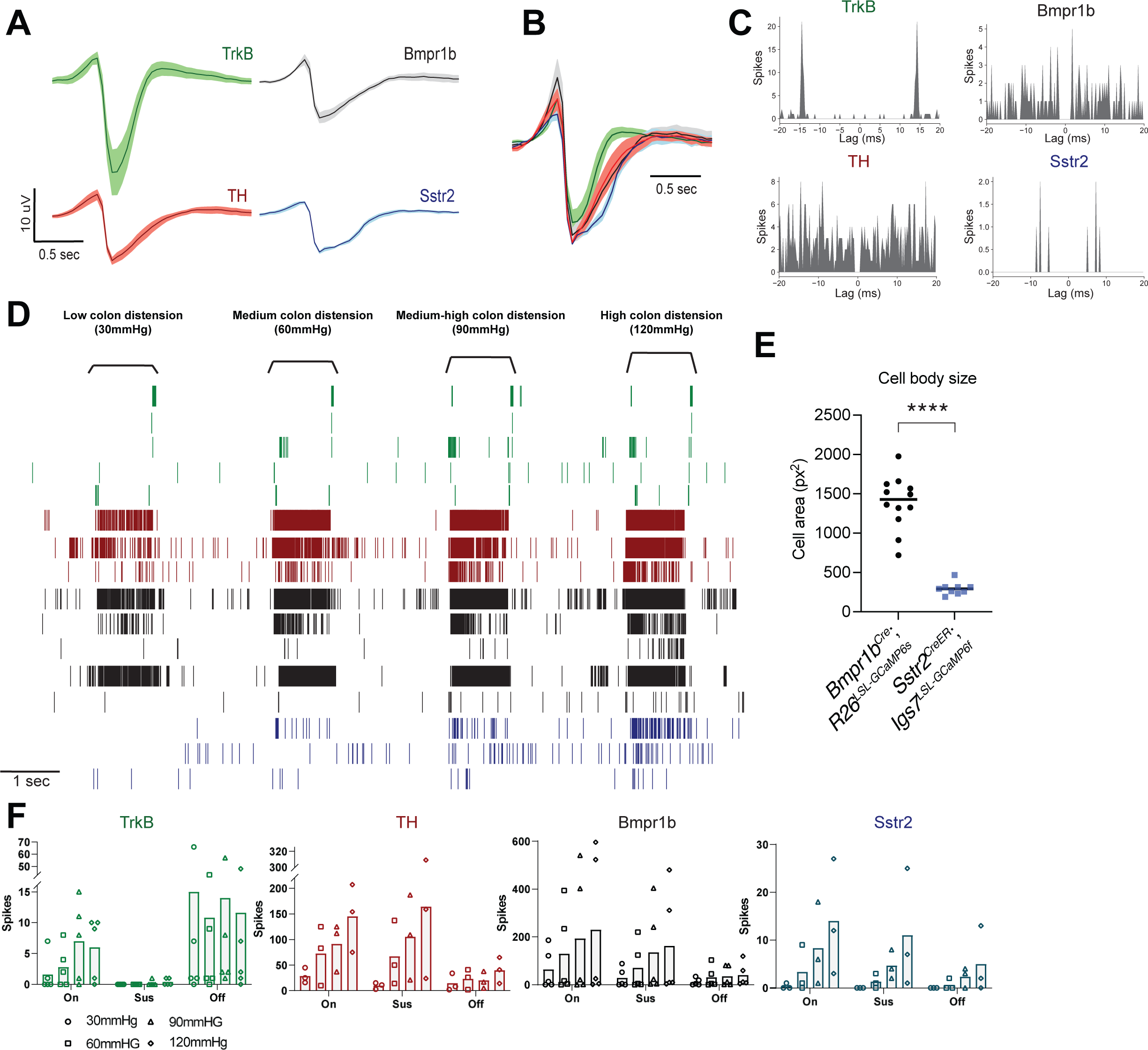
*In vivo* MEA physiological properties and firing rates in response to colon distension of four DRG afferent subtypes. **A**: Raw action potential (AP) waveforms of each DRG afferent subtype (mean +/-SEM). **B**: Normalized AP waveforms of each DRG afferent subtype (mean +/-SEM). **C**: Example autocorrelograms of each DRG afferent subtype. **D**: Cell size quantification of GCaMP-expressing cells from *Bmpr1b^Cre^*; *R26^LSL-GCaMP6s^* (n = 12 cells from N = 2 DRG) and *Sstr2^CreER^*; *Igs7^LSL-GCaMP6f^* (n = 9 cells from N = 1 DRG). Compared with an unpaired t test. **E**: Raster plot with all opto-tagged units, organized by genotype. TrkB^+^ units in green (n = 5 cells from N = 5 mice); TH^+^ units in red (n = 3 cells from N = 3 mice); Bmpr1b^+^ units in black (n= 5 cells from N = 3 mice); and Sstr2^+^ units in blue (n = 3 cells from N = 3 mice). **F**: Firing rates of each DRG afferent subtype during the on, sustained (sus), and off periods at each colonic distension force.

**Supplementary Figure 6:**
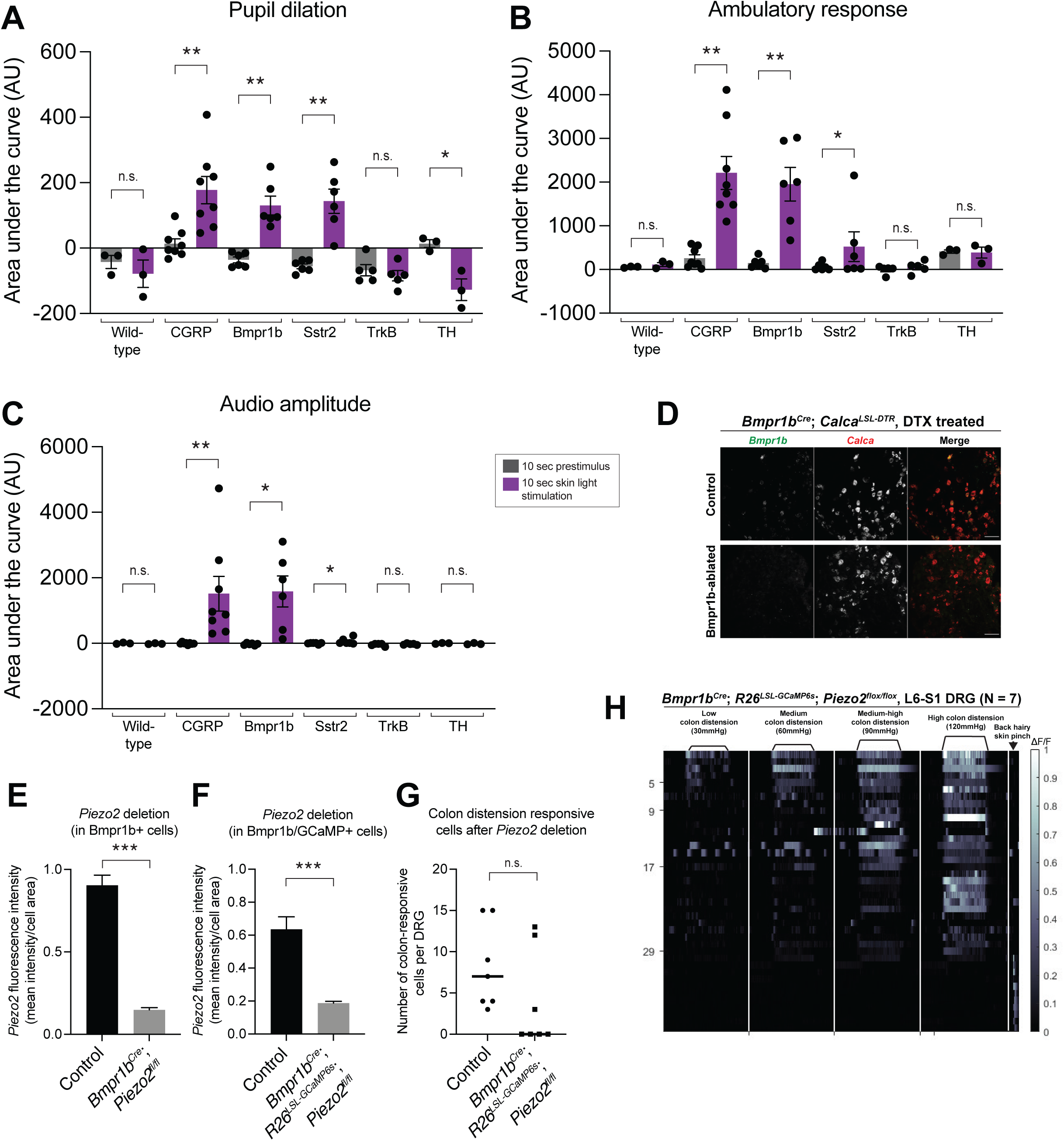
Piezo2 deletion does not abolish the response to high threshold colonic distension in Bmpr1b^+^ DRGs. **A-C**: Behavioral responses (pupil dilation (**A**), movement (**B**), and vocalizations (**C**)) to skin (back hindpaw and leg) optogenetic stimulation in wild-type mice (N = 3), *Phox2B-Flp; R26^FSF-^ mCitrine-ReaChR* _mice (N = 6), *Calca-Flp*; *R26*_*FSF-ReaChR* _or *Calca*_*CreER*_; *R26*_*FRT-LSL-ChR2-YFP-FRT* _mice (N_ = 8), Bmpr1b^Cre^; Calca-Flp; R26^FSF-LSL-mCitrine-ReaChR^ mice (N = 6), Sstr2^CreER^; Calca-Flp; R26^FSF-^ LSL-mCitrine-ReaChR _or Sstr2_CreER_; R26_LSL-mCitrine-ReaChR _mice (N = 6), TrkB_CreER_; Advillin_Flp_; R26_FSF-LSL-^mCitrine-ReaChR^ mice (N = 5), and TH^CreER^; Phox2B-Flp; R26^FRT-LSL-ChR2-YFP-FRT^ mice (N = 3). All comparisons are paired t tests except the accelerometer data for Sstr2^CreER^; Calca-Flp; R26^FSF-LSL-mCitrine-ReaChR^ mice and and TH^CreER^; Phox2B-Flp; R26^FRT-LSL-ChR2-YFP-FRT^ mice and vocalization data for Calca-Flp; R26^FSF-ReaChR^ or Calca^CreER^; R26^FRT-LSL-ChR2-YFP-FRT^ mice, Sstr2^CreER^; Calca-_Flp; R26_FSF-LSL-mCitrine-ReaChR _or Sstr2_CreER_; R26_LSL-mCitrine-ReaChR _mice, and TrkB_CreER_; Advillin_Flp_;_ R26^FSF-LSL-mCitrine-ReaChR^ mice, which were analyzed using the Wilcoxon matched-pairs signed rank test. **D**: DRG *in situ* hybridization in control and Bmpr1b-ablated animals. **E-F**: Quantification of Piezo2 deletion efficiency based on DRG *in situ* hybridization in control and *Bmpr1b^Cre^*; *Piezo2^fl/fl^* or *Bmpr1b^Cre^*; *R26^LSL-GCaMP6s^*; *Piezo2^fl/fl^* mice (comparisons are unpaired t tests). **G**: Number of colon distension responsive cells in control and *Bmpr1b^Cre^*; *R26^LSL-GCaMP6s^*; *Piezo2^fl/fl^* DRGs (analyzed using the Mann-Whitney test). Note that the control neuron measurements are data that is replotted from Figure 4. **H**: *In vivo* DRG calcium imaging responses to colon distension in *Bmpr1b^Cre^*; *R26^LSL-GCaMP6s^*; *Piezo2^fl/fl^* (N = 7) animals.

**Supplementary Figure 7:**
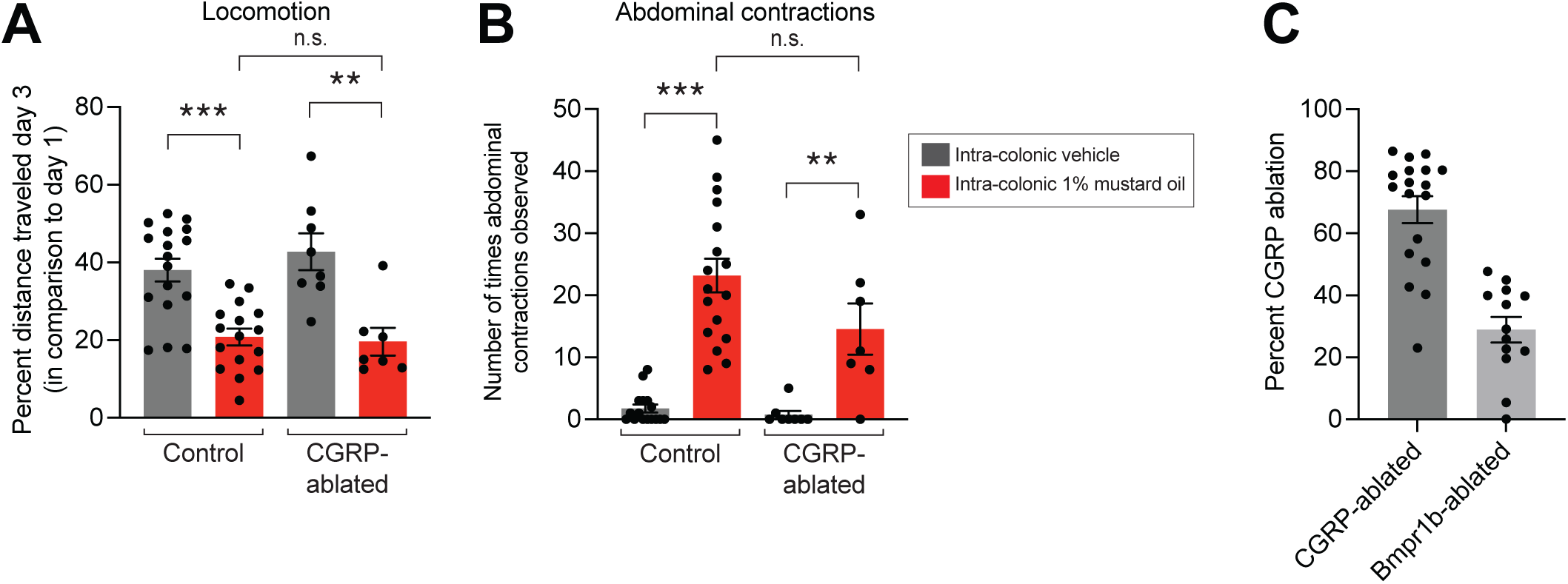
CGRP ablation does not affect the response to intracolonic mustard oil. **A-B**: Behavioral response to intracolonic mustard oil in control and CGRP-ablated (*Advillin^Cre^*; *Calca^LSL-hDTR^*) animals. All comparisons are unpaired t tests (N = 17 for controls, vehicle treated; N = 16 for controls, mustard oil treated; N = 8 for CGRP-ablated, vehicle treated; N = 7 for CGRP-ablated, mustard oil treated). **C**: Percent CGRP ablation in CGRP-ablated (*Advillin^Cre^*; *Calca^LSL-hDTR^*) and Bmpr1b-ablated (*Bmpr1b^Cre^*; *Calca^LSL-hDTR^*). Each point represents one animal, which was calculated from the average of 5 DRGs per animal. For CGRP-ablated experiments, only animals with >75% CGRP ablation were included in analyses (animals in Figure S7A-B). For Bmpr1b-ablated experiments, only animals > 22% CGRP ablation (correlated to > 75% Bmpr1b ablation) were included in analyses (animals in Figure 6D-F, 7A, E-G).

## Methods

### Key Resource Table

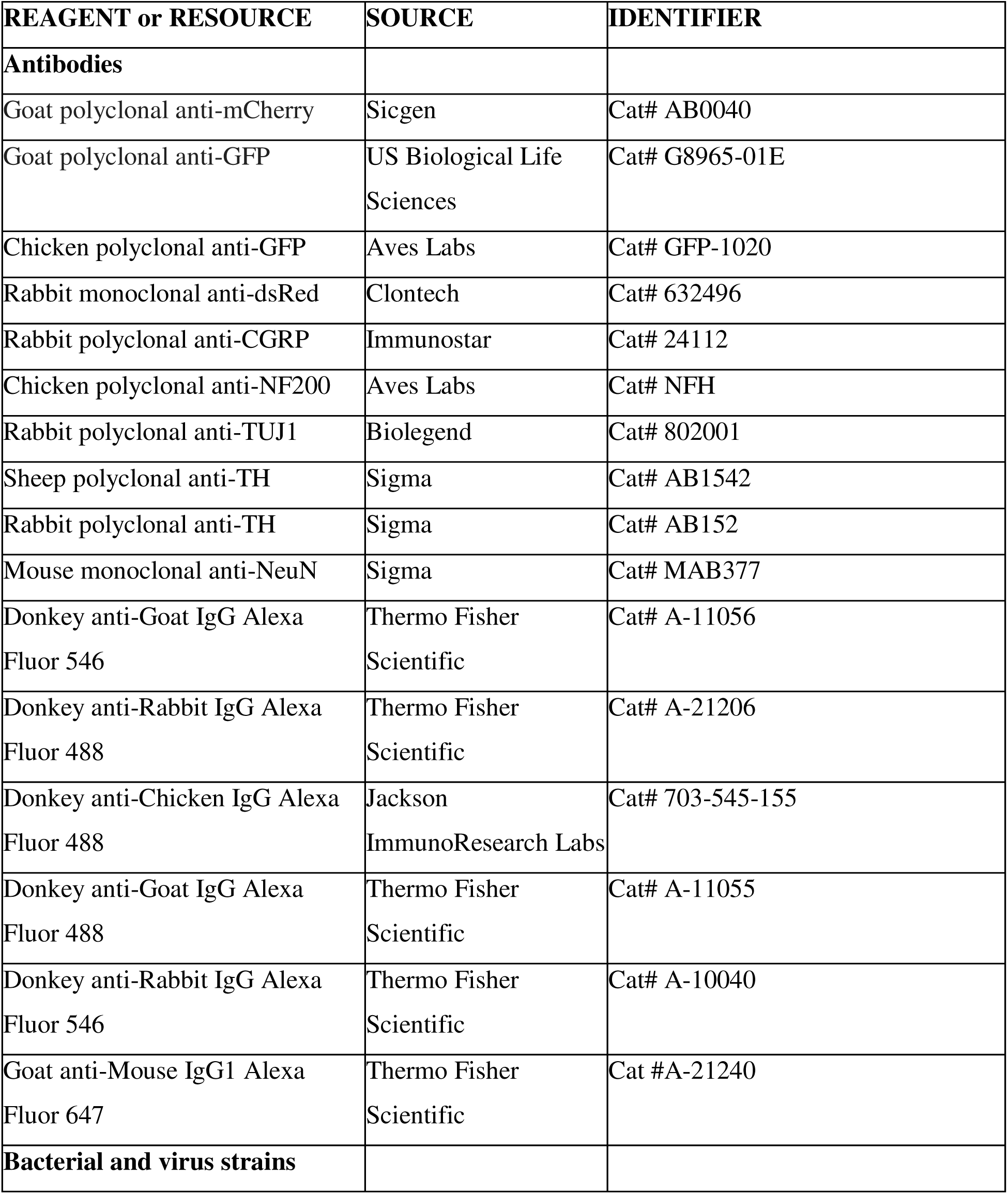
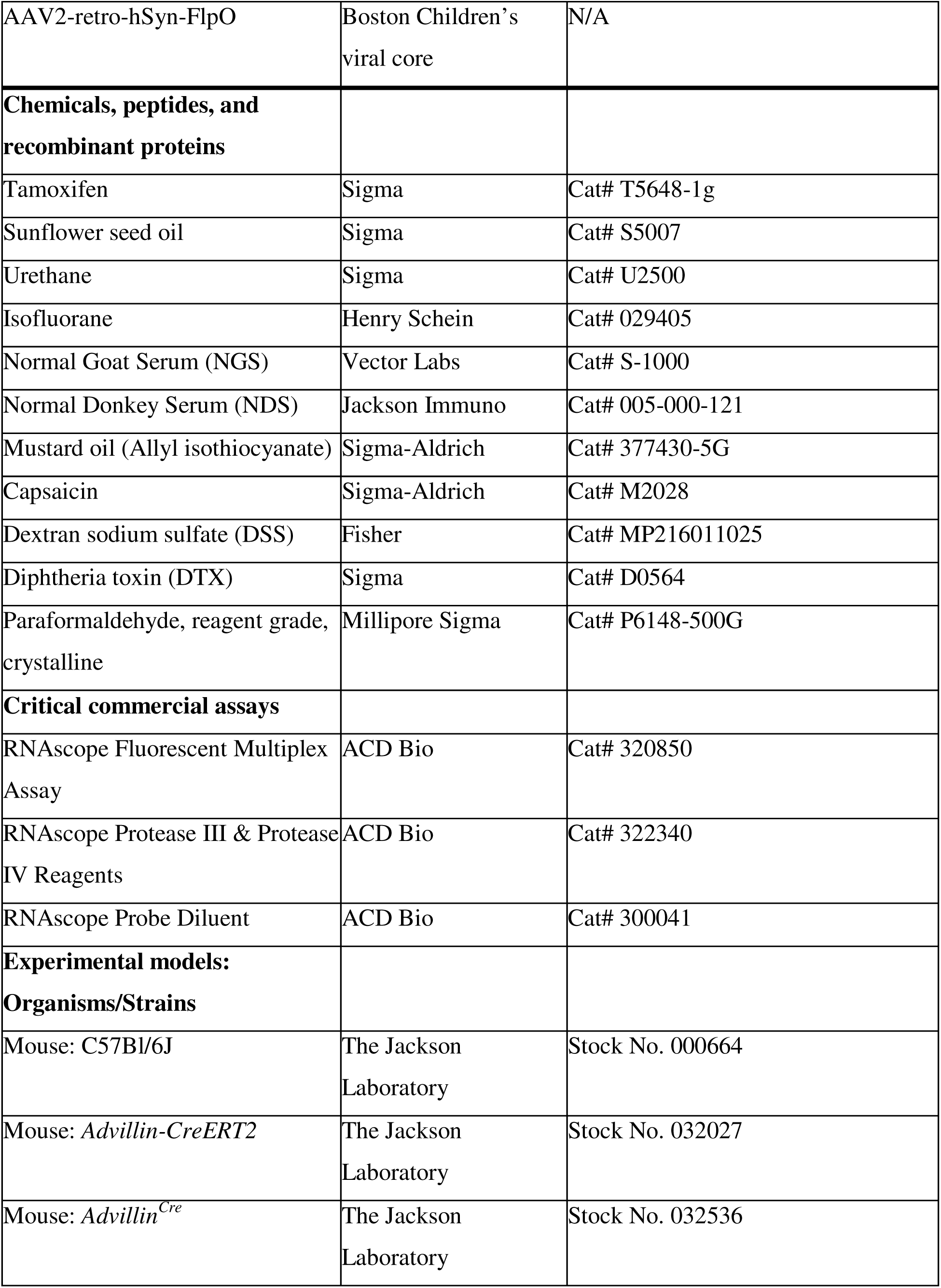
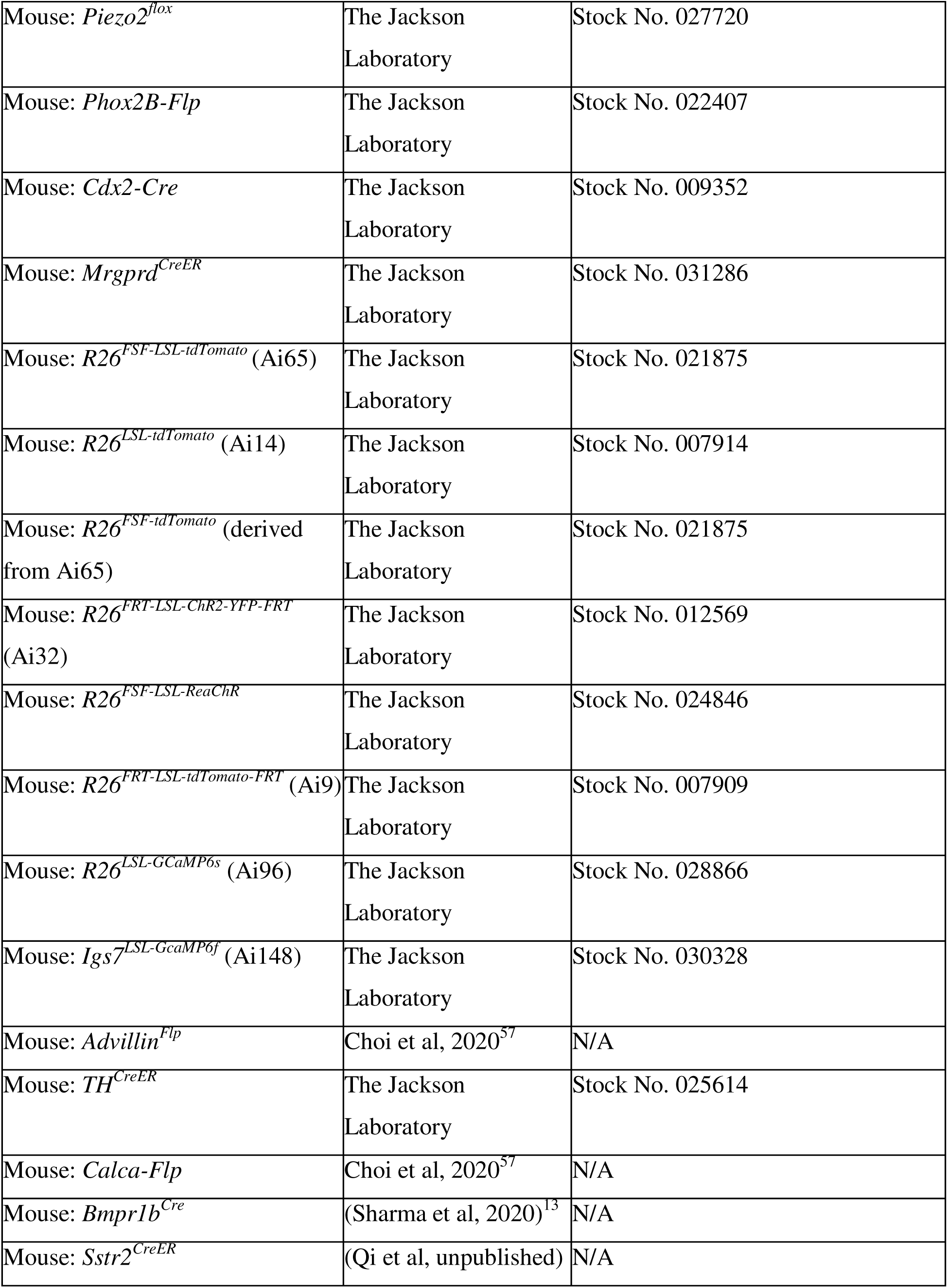
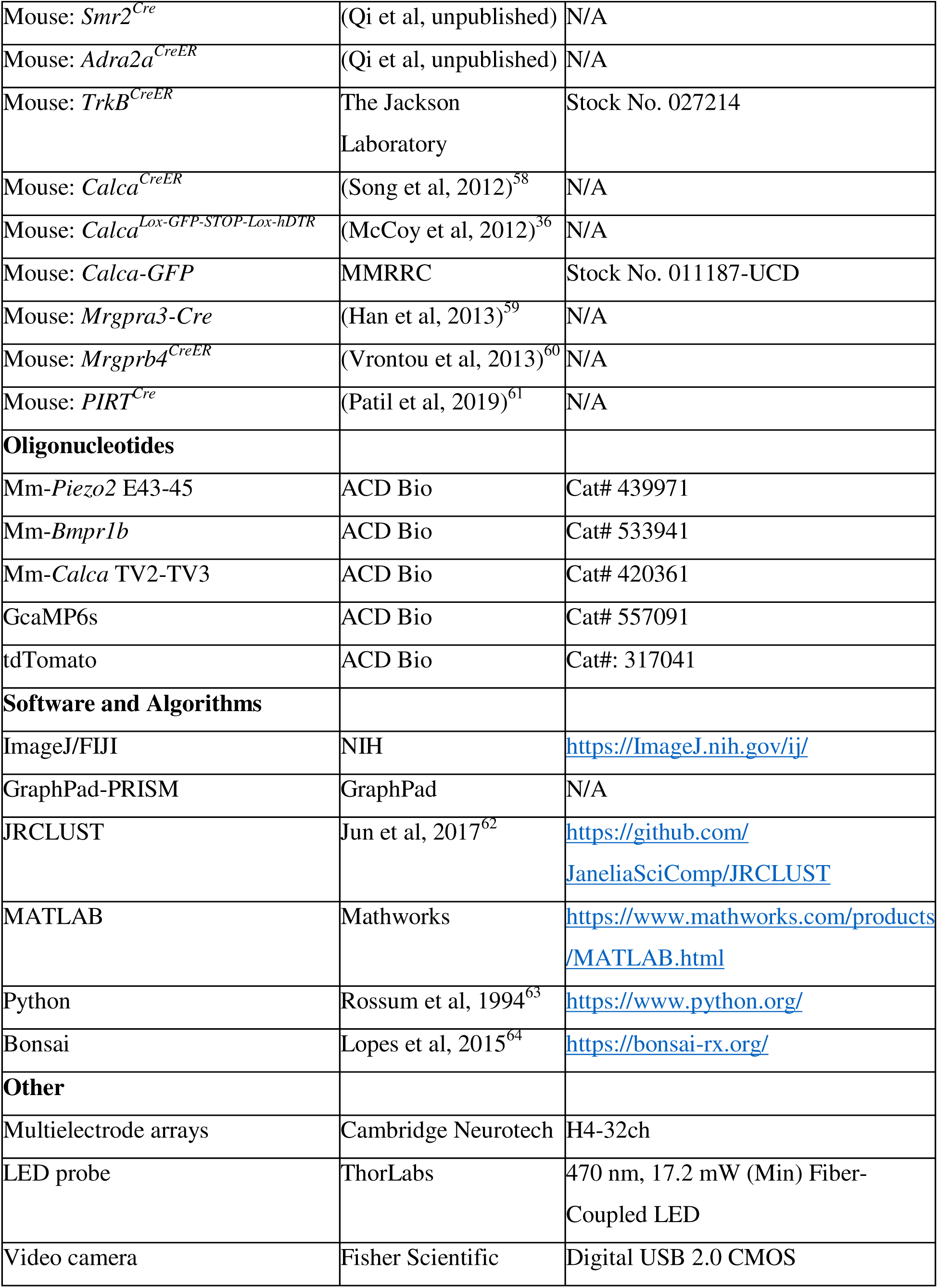
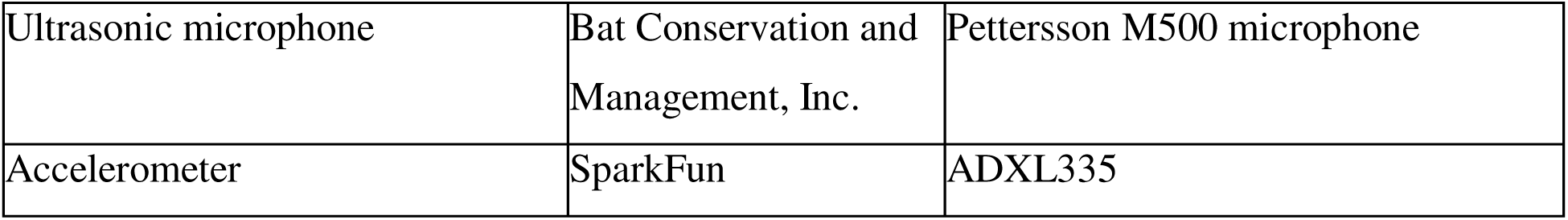

## Resource availability

### Lead contact

Further information and requests for resources and reagents should be directed to and will be fulfilled by the lead contact, David Ginty.

### Material availability

All new reagents will be made available upon request.

### Animals

All mouse experiments in this study were approved by the National Institutes of Health and the Harvard Medical School IACUC. Experiments followed the ethical guidelines outlined in the NIH Guide for the care and use of laboratory animals (https://grants.nih.gov/grants/olaw/guide-for-the-care-and-use-of-laboratory-animals.pdf). Mice were housed under standard conditions and given chow and water *ad libitum*. *Bmpr1b^T2a-CreER^* mice were generated using standard CRISPR-based homologous recombination techniques in embryonic stem (ES) cells. Chimeras were generated by blastocyst injection and subsequent germline transmission was confirmed by tail PCR. The neo selection cassette was excised using a Flp-deleter strain.

The following mice are available from Jax: *Advillin-CreERT2* (Stock No. 032027), *Advillin^Cre^* (Stock No. 032536), *Piezo2^flox^* (Stock No. 027720), *Phox2B-Flp* (Stock No. 022407), *Cdx2-Cre* (Stock No. 009352), *Mrgprd^CreER^* (Stock No. 031286), *R26^FSF-LSL-tdTomato^* (Ai65; Stock No. 021875), *R26^LSL-tdTomato^* (Stock No. 007914), *R26^FSF-tdTomato^* (derived from Ai65; Stock No. 021875), *R26^FRT-LSL-ChR2-YFP-FRT^* (Ai32; Stock No. 012569), *R26^FRT-LSL-tdTomato-FRT^* (Ai9; Stock No. 007909), *R26^FSF-LSL-ReaChR^* (Stock No. 024846), *R26^LSL-GcaMP6s^* (Ai96; Stock No. 028866), and *Igs7^LSL-GcaMP6f^* (Ai148; Stock No. 030328). Mice generated in the Ginty lab (and available at JAX now or in the near future) are: *Advillin^Flp^* (Choi et al, 2020)^57^, *TH^CreER^* (Stock No. 025614)^14^, *Calca-Flp* (Choi et al, 2020)^57^, *Bmpr1b^Cre^* (Sharma et al, 2020)^13^, *Sstr2^CreER^* (Qi et al, unpublished), *Smr2^Cre^* (Qi et al, unpublished), *Adra2a^CreER^* (Qi et al, unpublished), and *TrkB^CreER^* (Rutlin et al, 2014)^7^. Other mice used in the study have been previously described: *Calca^CreER^* (Song et al, 2012)^58^, *Calca^Lox-GFP-STOP-Lox-hDTR^* (McCoy et al, 2012)^36^, *Calca-GFP* (MMRRC, Stock No. 011187-UCD), *Mrgpra3-Cre* (Han et al, 2013)^59^, *Mrgprb4^CreER^* (Vrontou et al, 2013)^60^, and *PIRT^Cre^* (Patil et al, 2019)^61^.

All experiments with wild-type animals were conducted with mice on the C57Bl/6J background obtained from Jackson Laboratory.

### Tamoxifen treatment

Tamoxifen was dissolved in ethanol (20 mg/ml). It was then mixed with an equal volume of sunflower seed oil (Sigma) and vortexed for 20 minutes. The mixture was subsequently centrifuged under vacuum for 30 minutes to remove the ethanol. The solution was kept at −80u. It was delivered by oral gavage to pregnant females for embryonic treatment, or by intraperitoneal (IP) injection for postnatal treatment. For all analyses, the plug date was designated as E0.5 and the date of birth as P0.

Mice were treated with tamoxifen at the following timepoints (by genotype) unless otherwise specified: *Advillin-CreER* (P21-25); *Mrgprd^CreER^* (P6-P9); *TrkB^CreER^* (P3-5); *TH^CreER^* (P21); *Calca^CreER^* (P11); *Sstr2^CreER^* (P21); *Adra2^CreER^* (P14 and P16); *Bmpr1b^CreER^* (P21).

### Whole mount immunofluorescence

The whole-mount immunofluorescence (IF) was performed as previously described^7^ for skin samples. The protocol was adapted for optimized use in the GI tract. Mice (P21-42) were euthanized with isoflurane and perfused with 5ml PBS followed by 10-20 ml of 4% paraformaldehyde (PFA) in PBS at room temperature (RT). Skin and the GI tract were dissected from perfused mice. Hair was removed from the skin using commercial depilatory cream (NAIR, Church and Dwight Co.; Princeton, NJ) for 0.5-1 min and washed with water. The colon was dissected open and the luminal contents were removed. Skin and colon were post-fixed in Zamboni’s fixation buffer at 4°C for 12-24 hours. Thereafter, the samples were washed with PBS for 3×5 minutes, followed by PBST (PBS with 1% Triton X-100) every 20 minutes for 5-7 hours. A small piece of each tissue was cut (0.5mm x 0.5mm) and incubated in primary antibodies diluted in blocking solution (75% 1% PBST, 20% DMSO, 5% Normal Goat Serum (NGS) (Vector Labs, S-1000) or Normal Donkey Serum (NDS) (Jackson Immuno, 005-000-121)) for 3-4 days on a rocking platform at RT. The samples were thereafter washed again in PBST for 5-7 hours every 20 minutes and then incubated with secondary antibodies in blocking solution on rocking platform for 2 days at RT. The samples were again washed in PBST for 5-7 hours every 20 minutes. The samples were subsequently dehydrated: 1 wash for 10 minutes in 50% methanol (MeOH), 1 wash for 10 minutes in 80% MeOH, followed by 3 washes for 20 minutes in 100% MeOH. The samples were stored in 100% MeOH at −20°C. To image, the samples were cleared in BABB (1 part Benzyl Alcohol: 2 parts Benzyl Benzoate) for 5 minutes, and then mounted on slides using BABB as the mounting medium.

### Tissue section immunofluorescence

Mice were perfused as above. Spinal columns containing spinal cords, DRGs, and sympathetic chain were dissected, post-fixed at 4°C in a PFA (4% in PBS) solution for 12-24 hours, and subsequently washed 3×5 minutes in PBS. For DRG and spinal cord IF, the spinal cord and DRGs were dissected out after fixation. For DRG, spinal cord, and sympathetic chain IF, the spinal column was left intact. Tissues were cryoprotected in 30% sucrose in PBS at 4°C for two nights and thereafter embedded in OCT and frozen at -20°C. Tissues were sectioned at 20-30μm with a cryostat. Slides were rehydrated with PBS 3×5 minutes and then blocked with PBS containing 0.1% Triton X-100 and 5% NGS or NDS for 20-60 minutes at room temperature. Tissue sections were incubated with primary antibodies diluted in PBS with 5% NGS or NDS overnight at 4°C. The following day, sections were washed 4×5 minutes in PBS containing 0.02% Triton X-100 and then incubated with secondary antibodies diluted 1:500 in PBS with 5% NGS or NDS for 1 hour at RT. The sections were washed again 4×5 minutes in PBS containing 0.02% Triton X-100 and mounted with fluoromount-G (Southern Biotech).

Primary antibodies used for both whole mount and tissue section IF include: chicken anti-GFP (GFP-1020, Aves Labs; 1:1000), goat anti-GFP (G8965-01E; US Biological Life Sciences; 1:500), rabbit anti-DsRed (632496; Clontech; 1:500), goat anti-mCherry (AB0040-200, Acris, 1:500), rabbit anti-CGRP (Immunostar, 24112, 1:500), chicken anti-NF200 (Aves Labs, NFH, 1:1000), rabbit anti-TUJ1 (Biolegend, 802001, 1:500), sheep anti-TH (Sigma, AB1542; 1:500), rabbit anti-TH (Sigma; AB152; 1:500), and mouse anti-NeuN (Sigma; MAB377; 1:500).

Secondary antibodies used for both whole mount and tissue section IF included: Alexa 488, 546 or 647 conjugated goat anti-chicken antibodies, Alexa 488, 546 or 647 conjugated goat anti-rabbit antibodies, Alexa 488, 546 or 647 conjugated donkey anti-goat antibodies, Alexa 488, 546 or 647 conjugated donkey anti-rabbit antibodies, Alexa 488, 546 or 647 conjugated donkey anti-goat antibodies. All secondary antibodies above were purchased from Life Technologies, except Alexa 488/647 conjugated donkey anti-chicken antibodies, which were purchased from Jackson ImmunoResearch.

For all anatomy experiments, data was collected from at least N = 2 animals. All scale bars represent 100 μ unless otherwise specified.

### *In situ* hybridization

RNA fluorescence *in situ* hybridization (RNA-FISH) was performed as previously described^13^. Briefly, individual DRGs were rapidly dissected from mice and subsequently frozen in dry-ice-cooled 2-metylbutane and stored at −80°C until they were sectioned. DRG were sectioned at a thickness of 20 μ and RNAs were detected with RNAscope (Advanced Cell Diagnostics) using the manufacturer’s protocol. The following probes were used: Mm-*Bmpr1b* (Cat#: 533941), Mm-*Calca* TV2-TV3 (Cat#-420361), Mm-*Piezo2* E43-45 (Cat#: 439971), GcaMP6s (Cat#: 557091), and tdTomato (Cat#: 317041).

### Tissue processing for H&E

Animals were perfused and colons were dissected as above, although the colons were not dissected open, and their luminal contents were left intact. Colons were post-fixed at 4°C in 4% PFA for 12-24 hours, and subsequently washed 3×5 minutes in PBS. Tissues were embedded in paraffin and sectioned. Thereafter, slides were stained with standard hematoxylin and eosin (H&E) staining techniques and imaged on a light microscope (Olympus BX63).

### Single neuron morphological analysis

For sparse labeling of single neurons, mice were treated with approximately 1/10^th^ or 1/100^th^ the usual tam dose for each subtype. The tissues were processed by whole mount immunofluorescence. The endings were traced in ImageJ using Single Neurite Tracer (SNT). Parameters such as process length, width, and number of branches were calculated using SNT.

### Bmpr1b^+^ and CGRP^+^ neuron ablation analysis

To assess the percent of Bmpr1b^+^ neurons ablated per DRG in *Bmpr1b^Cre^*; *Calca^LSL-hDTR^* DRGs, we first used *in situ* hybridization to confirm in two animals that > 75% of the *Bmpr1b*^+^ cells were ablated. We also calculated the average percent of *Calca*^+^ cells that were also *Bmpr1b*^+^ per DRG in controls (average across N = 3 DRGs). For most animals, we used immunofluorescence to quantify the number of CGRP^+^ cells over the total number of cells per DRG (total number of cells quantified by NeuN staining). The average percent of *Bmpr1b*^+^ cells out of the total *Calca*^+^ population was then used to calculate the amount of Bmpr1b^+^ cell ablation per DRG. This was compared between controls and ablated animals.

To assess the percent of CGRP^+^ cells ablated per DRG in *Advillin^Cre^*; *Calca^LSL-hDTR^* DRGs, immunofluorescence was used to quantify the number of CGRP^+^ cells over the total number of cells per DRG (total number of cells quantified by NeuN staining). This was compared between ablated animals and an average of multiple controls to calculate percent ablation.

### *Piezo2* deletion quantification

To quantify *Piezo2* deletion in *Bmpr1b^Cre^*; *Piezo2^fl/fl^* or *Bmpr1b^Cre^*; *R26^GcaMP6S^*; *Piezo2^fl/fl^* animals, DRGs were processed for *in situ* hybridization. All experiments had control and knockout slides processed on the same day, with the same antibody conditions. Samples were also imaged on the same day using the same microscope settings, and all processing of the images was consistent between control and knock-out slides. For *Bmpr1b^Cre^*; *Piezo2^fl/fl^* DRGs, regions of interest (ROIs) were drawn in ImageJ around *Bmpr1b* positive cells. The total fluorescence intensity in the *Piezo2* channel of each of these regions was calculated in ImageJ and divided by the cell area to quantify the *Piezo2* expression per genotype. For the *Bmpr1b^Cre^*; *R26^GcaMP6S^*; *Piezo2^fl/fl^* DRGs, the ROIs were drawn around the GcaMP6S cells and processed in an analogous fashion.

### Cell size quantification

To quantify cell size in *Bmpr1b^Cre^*; *R26^LSL-GcaMP6s^* and *Sstr2^CreER^*; *Igs7^LSL-GcaMP6f^* mice, DRGs were processed for *in situ* hybridization. Samples were imaged with the same microscope settings and all processing of the images was consistent between genotypes. ROIs were drawn in ImageJ around GcaMP^+^ cells and the area (in px^2^) of each ROI was calculated using ImageJ.

### *In vivo* calcium imaging and electrophysiology

Mice were anesthetized with urethane (0.5-1.5 mg/g) and placed on a bite bar (Kopf Instruments) on a custom heated recording platform. Temperature was monitored throughout the procedure with a temperature controller (TC-344B, Warner Instruments) and maintained at 35.5-37.5°C with a thermoelectric heater (C3200-6145, Honeywell) embedded in castable cement (Aremco). To expose L6-S1 DRGs, an incision was made over the spine (T10-S2) and the overlying tissue was retracted, exposing the vertebral canal. The vertebral canal was secured with custom spinal clamps (made in the Harvard Medical School Neurobiology Department Machine Shop) and the bone dorsal to the target DRG was removed using forceps.

The exposed L6-S1 DRG was then used for calcium imaging with a Zeiss Axio Examiner microscope or multiunit electrophysiological recordings using an MEA probe (H4-32ch, Cambridge Neurotech) that was inserted into the DRG for recording.

A balloon attached to a manometer (Cole Parmer) was inserted into the colon of the mice. For all calcium imaging and electrophysiology experiments, each colon distension measurement involved a 1-minute baseline, 1 minute with the balloon distended to the specified pressure, and a 1-minute period after deflation for a total 3-minute recording. For hairy skin pinch measurements, the pinch was performed using forceps after a 10-second baseline and with a 10 second period after stimulus, for a total 20-second recording.

For colon optogenetic stimulation in MEA experiments, the MEA probe was left in place in the DRG and the colon balloon was replaced with an LED probe (ThorLabs; 470 nm, 17.2 mW (Min) Fiber-Coupled LED) inserted 1cm into the colon. Opto-responsive units were determined by delivering brief (5-20ms) pulses of blue light (insert power) to the colon. The first spike to each light pulse was used to determine the optogenetic latency.

### Spike sorting

Spike sorting was performed as previously described^65,66^. Briefly, open-source software (JRCLUST version 3.2.2) was used to automatically sort action potentials into clusters, manually refine clusters, and classify clusters as single or multi units. The voltage traces were filtered with a differentiation filter of order 3. Frequency outliers were removed with a threshold of 10 median absolute deviations (MADs). Action potentials were detected with a threshold of 4.5 times the standard deviation of the noise. Action potentials with similar times across sites within 60 mm were merged and action potentials were then sorted into clusters with a density-based-clustering algorithm (clustering by fast search and find of density peaks) with cutoffs for log_10_® at -3 and log_10_(d) at 0.6. Clusters with a waveform correlation greater than 0.99 were automatically merged and outlier spikes (> 6.5 MADs) were removed from each cluster.

Manual cluster curation was performed with JRCLUST split and merge tools. Single units were classified based on waveform consistency, spike amplitude, clear refectory periods (interspike intervals of > 1ms), and waveform separability with nearby clusters. Only clusters determined to be single units were used. Spike times for single units were exported and analyzed in Python (3.8.5).

### Calcium imaging analysis

The motion correction of the calcium imaging videos was performed in ImageJ using the moco plugin^67^. Individual responding ROIs were manually segmented. Measured intensity for each ROI was further processed using a custom MATLAB code. Response was defined as 2.5 times over the standard deviation of the baseline. In the heatmap plot, cells of the same subtype were sorted by the maximum response.

### Dorsal horn injections

Mice (P21–P28) were anesthetized by continuous inhalation of isoflurane (1.5–2.5%) using an isoflurane vaporizer (VetEquip). Laminectomies were performed to expose the lumbar spinal cord at L6, and a total of 450 nl of AAV virus (AAV2-retro-hSyn-FlpO, Boston Children’s Core; titer 2.06×10^14^ gc/ml) was directly injected into two or three adjacent spots in the dorsal spinal cord using pulled glass pipettes (Wiretrol II, Drummond) and a microsyringe pump injector (UMP3, World Precision Instruments).

### Behavioral testing

Male and female mice of mixed genetic background were used for all behavior experiments. Testing was done with mice between 6-12 weeks of age. All animals were group housed and controls for most experiments were littermates. The exceptions to this were three of the controls in the *Cdx2-Cre*; *Piezo2^flox/flox^* balloon distension experiments and wild-type optogenetic experiments, for which wild-type C57BL/6J animals were used. For all behavioral assays, mice were habituated to the behavior room in their home cage for 20 minutes each day, prior to either a habituation or test day.

### Headplating

For all behavior experiments in which pupil dilation, movement, and vocalizations were recorded, the mice were headplated. For this, mice were anaesthetized with 1.5–2% isoflurane, the scalp was removed, the skull was dried, and a titanium headplate was affixed to the skull using dental cement (Metabond). The mice were treated with sustained release buprenorphine (0.1 mg/kg) after the procedure. Mice were allowed at least 5-7 days to recover after the procedure prior to undergoing behavioral testing.

### Colon distension and optogenetic behavioral testing

The behavioral setup for colon distension and optogenetic testing involved a bar to which the headplate could attach mounted next to a custom tube, to body fix the animals. The tube was mounted on foam to allow some vibration with movement of the mouse. The accelerometer was attached to the outside of the tube. A video camera (digital USB 2.0 CMOS) was mounted to record the eye and a microphone which could record in the ultrasonic range (Bat Conservation and Management, Inc.) was mounted at the front of the setup. The setup was placed in a sound-attenuating box (Med. Associates, Inc.) to decrease ambient noise.

For habituation, the mice were head-fixed and body-fixed in the setup for 5 minutes on the first day and 10 minutes on the second day. An investigator held their tail in place to re-capitulate the day of the assay. The behavioral test was performed after 2 days of habituation.

For colon distension testing, the intracolonic balloon was built as previously described^68^. The balloon, which was connected to a manometer (Cole Parmer) was inserted into the colon of the mice and secured to the tail with tape. An investigator lightly held the tail in place to avoid balloon movement causing damage to the colon. Each distension trial was 5 minutes in total and included a 1-minute baseline, followed by four balloon inflations (lasting 10 seconds each) at each minute mark. The balloon was inflated to 2.4PSI within 2 seconds and held at 2.4PSI for the 10 second stimulus duration. Each animal was tested for at least 1 trial. If an animal displayed stressful behaviors such as kicking during the baseline period, then the trial was repeated and the trial with the least amount of baseline stress was used for the final analysis.

For colon optogenetic testing, an LED was inserted 1cm into the colon and secured to the tail with tape. An investigator lightly held the tail in place to avoid LED movement causing damage to the colon. Each trial was 5 minutes in total, with a 1-minute baseline, followed by four 10-second blue light stimulations at 10Hz. For animals expressing Channelrhodopsin2 (ChR2), 60mW power was used, while for animals expressing Red-shifted variant of Channelrhodopsin (ReaChR), 36mW power was used. For each animal, the optogenetic stimulation was repeated three times and the average of the three trials was used for each animal. After colon optogenetic activation, the LED was removed from the colon, cleaned, and optogenetic stimulation of the back left paw was performed with the same experimental paradigm as colon stimulation.

The blue LED (Doric Lenses) was connected to a programmable LED driver (Doric Lenses). Optogenetic stimulation was controlled by the combination of custom programs written in Bonsai software and Doric Neuroscience Studio (4.1.5.2) through an Arduino circuit board (Uno, Arduino) and custom sketches written in Arduino software (1.8.7).

### Pupil dilation, movement, and vocalization analysis

The pupil dilation analysis was performed as previously described^57^. Briefly, pupils were recorded at 30 frames s^−1^ using a digital USB 2.0 CMOS video camera mounted near one eye. An infrared illuminator was used to obtain clear images of the pupil. The pupil diameter was tracked at 10Hz using a custom program written in Bonsai software.

Movement (ambulatory response) data was collected using an accelerometer (SparkFun ADXL335) connected to a data acquisition card (National Instruments). The movement in the X, Y, and Z axes was summed at each time point for the total movement. The data were collected at 20Hz using a custom program written in Bonsai software and analyzed using a custom program written in Matlab.

Vocalizations (including in the ultrasonic range) were recorded using one of two microphones (UltraVox or Bat Conservation and Management, Inc.). The amplitude and frequency of sound were recorded at 20Hz using a custom program written in Bonsai software and analyzed using a custom program written in Matlab.

The average baseline pupil diameter, ambulatory response, or audio amplitude was the average of those values during the 1-minute baseline period before the beginning of the first optogenetic stimulation or balloon distension. The value at each time point was divided by the average of the 1-minute baseline to obtain a normalized value. This value was subtracted from 1 and multiplied by 100 to obtain a percent change in pupil dilation, movement, or vocalizations. The area under the curve (AUC) in the 10 second prestimulus time period and 10 second stimulus period were calculated using a custom program written in Matlab for each stimulation. The average prestimulus and stimulus values were calculated for each mouse (across either four balloon distensions in one five minute trial for the balloon assay, or 12 total optogenetic stimulations across three trials for the optogenetic activation assay).

### Diphtheria toxin (DTX) mediated ablation

*Bmpr1b^Cre^*; *Calca^LSL-hDTR^* or *Advillin^Cre^*; *Calca^LSL-hDTR^* mice were treated every day for 21 days with intraperitoneal (IP) injections of DTX (0.175 μg total over 21 injections) starting at age 6 weeks. Animals were tested in behavioral assays 18 days after completing DTX injections. Ablation efficiency was calculated as described above. Data shown are from animals with > 75% ablation.

### Colonic mustard oil behavioral testing

Colonic mustard oil behavioral testing was performed as previously described^69^. Briefly, on day 1, animals were habituated for 20 minutes in the chambers, which were 10cm x 10cm x 10cm acrylic chambers with one clear wall and three black walls. During this 20-minute period on day 1, animals were tracked with an overhead camera to record their baseline movement in the open field assay. On day 2, animals were again habituated for 20 minutes to the chamber.

The behavioral test was performed on day 3. Prior to the start of the assay, animals were anesthetized briefly with isoflurane. 50μl of either vehicle (70% ethanol dissolved in sterile saline) or 1% mustard oil in vehicle (Sigma) was inserted ∼1cm into the distal colon using a 24G reusable oral gavage needle (Fine Science Tools (FST)). Animals were then placed into the chamber and given 5 minutes to recover from the brief anesthesia. Their movement on an overhead camera as well as their behaviors on a camera mounted in front of the test chamber were recorded for 20 minutes.

For the distance traveled analysis, the mouse centroid was tracked, and speed and velocity of mouse locomotion were analyzed at 2 Hz with video files taken using an overhead camera and a custom program written in Bonsai software^57^. The distance traveled on day 3 (the day of the assay) was normalized to distance traveled at baseline (day 1) for each animal. Abdominal licking and contractions were analyzed manually using video files filmed by the camera mounted in front of the test chambers by investigators who were blinded to genotype.

### Dextran sodium sulfate (DSS)-induced colitis

To induce colitis, mice were treated with a solution containing dextran sodium sulfate (DSS; 4-5%) (MP Biomedical) dissolved in the drinking water. Controls were allowed access to regular drinking water. Animals were weighed on day 0 (the day when DSS was added to the drinking water). On each day of DSS treatment, weight and stool characteristics were recorded for each animal. This allowed for monitoring of the disease activity index (DAI), as previously described^70^. Animals were allowed access to 4-5% DSS in the drinking water for 6 days, followed by 1 day of regular drinking water. The behavioral and physiological experiments were performed on day 7, and the animals were perfused immediately thereafter for histological analysis.

### Statistical analyses

Statistical analyses were performed using GraphPad Prism (Version 8, GraphPad Software). The number of mice and the statistical tests used for individual experiments are included in the figure legends. Both non-parametric tests and parametric tests were used, depending on data normality, for comparing two independent groups (Mann-Whitney U test or Student’s t test) or two related groups (Wilcoxon matched-pairs signed rank test or paired Student’s t test). The following symbols are used in the figure legends for P values: n.s., not significant; *, P < 0.05; **, P < 0.01; ***, P < 0.001; ****, P < 0.0001. For all analyses, power analyses were conducted to calculate the ideal sample size for each experiment following experiments using 3-5 mice in each behavioral assay. Mice were randomly allocated into experimental groups when possible. All graphs show mean +/-standard error of the mean (SEM). For histological analyses, images were analyzed by investigators who were blinded to genotype whenever possible. For the behavioral assays, approximately half of the initial experiments were performed by experimenters who were not blinded to genotype, and the remainder of the experiments were performed by experimenters who were blinded to genotype, when possible. The data for all behavioral experiments were analyzed by experimenters who were blinded to genotype. Blinding was not used for the collection of data in electrophysiological recordings, but the data were analyzed automatically using Matlab or Python with the same scripts run for each experimental group.

## Acknowledgements

We thank P. Gorelik (HMS Research Instrumentation Core) and M. DeLisle for technical assistance, and Ginty lab members for discussion and comments on the manuscript. We also thank X. Dong for the *PIRT^Cre^* mice. This work was supported by NIH grants T32-GM007753 and 1-R25-AI147393-01 (R.L.W), DP2 NS127278 (N.S.), and NS097344 and AT011447 (D.D.G.), NEI P30 Core Grant for Vision Research EY012196 (O.M.), Klingenstein-Simons Foundation (N.S.), Whitehall Foundation (N.S.), The Hock E. Tan and Lisa Yang Center for Autism Research (D.D.G.), and the Lefler Center for Neurodegenerative Disorders (D.D.G.). D.D.G. is an HHMI investigator. This article is subject to HHMI’s Open Access to Publications policy. HHMI lab heads have previously granted a nonexclusive CC BY 4.0 license to the public and a sublicensable license to HHMI in their research articles. Pursuant to those licenses, the author-accepted manuscript of this article can be made freely available under a CC BY 4.0 license immediately upon publication.

## Author contributions

R.L.W. and D.D.G. conceived the study. R.L.W., A.A., and S.K. performed anatomical and behavior experiments. R.L.W. performed calcium imaging experiments, with input from L.Q. R.L.W. and G.R. performed *in vivo* DRG electrophysiological experiments. R.L.W., A.A., L.Q., and N.S. characterized mouse lines. R.L.W. analyzed data with assistance from L.Q., O.M., and S.C.; G.R. analyzed MEA data. R.L.W. and D.D.G. wrote the paper with input from all authors.

## Inclusion and diversity

We support inclusive, diverse, and equitable conduct of research.

